# Structural basis for the synergistic assembly of the snRNA export complex

**DOI:** 10.1101/2024.11.28.625805

**Authors:** Etienne Dubiez, William Garland, Maja Finderup Brask, Elisabetta Boeri Erba, Torben Heick Jensen, Jan Kadlec, Stephen Cusack

## Abstract

The nuclear cap-binding complex (CBC) and its partner ARS2 play crucial roles in regulating Pol II transcript fate through mutually exclusive interactions with RNA effectors. One such effector, PHAX, mediates the nuclear export of U-rich small nuclear RNAs (snRNAs). Here we present the cryo-EM structure of the snRNA export complex comprising phosphorylated PHAX, CBC, CRM1, Ran-GTP and capped RNA. The central region of PHAX bridges the CBC-bound capped RNA to the CRM1-RanGTP, while also significantly reinforcing cap dinucleotide binding. Additionally, PHAX interacts with a distant region of CRM1 facilitating contacts of an essential phosphorylated region with the prominent basic surface of RanGTP. The importance of these interactions is confirmed by *in vitro* and in cell mutagenesis experiments. CBC engagement within the snRNA export complex is incompatible with its interactions with other RNA effectors such as ALYREF or NCBP3. Taken together, we demonstrate that snRNA export complex formation requires synergistic binding of all its components, which in turn displaces ARS2 from the CBC, and commits the complex for export.

## Introduction

Eukaryotic transcription events generate vast amounts of RNA^1,2^. While useful transcripts are processed and mainly exported to the cytoplasm, the majority of RNAs are degraded in the nucleus^3^. Nascent RNA polymerase II (Pol II) transcripts acquire co-transcriptionally an m^7^G-cap structure at their 5’ end, which is subsequently bound by the nuclear cap-binding complex (CBC) composed of CBP80 and CBP20 (also known as NCBP1 and NCBP2)^4,5^. The cap is recognised by CBP20, while CBP80 interacts co-transcriptionally with numerous factors^5–10^. The CBC plays a central role in various steps of gene expression, including transcript synthesis and processing, nuclear export and nuclear RNA decay^5^.

During early Pol II transcription, the CBC interacts with the C-terminus of the Arsenite- Resistance Protein 2 (ARS2), which binds in a groove formed at the CBP20-CBP80 interface^9^. The CBC-ARS2 (CBCA) complex is a key regulator of RNA biogenesis, influencing transcriptional and post-transcriptional fate decisions for both productive and non-productive outcomes^11^. It is believed that the decision whether transcripts undergo processing, nuclear export or degradation depends on dynamic CBCA interactions with competing, mutually exclusive RNA effectors^7,11,12^. These effectors, which include the Phosphorylated Adaptor for RNA Export (PHAX), NCBP3 and ZC3H18, bind ARS2 via their ARS2-recognition motifs (ARMs) and the CBC through tryptophan-containing helices^7,13–15^. How such binding orchestrates the competitive interplay of RNA effectors remains poorly understood.

Correct RNA sorting is particularly crucial for short capped Pol II transcripts, as functionally important RNAs (e.g. snRNAs, snoRNAs and some histone and microRNAs) must be positively selected amidst a vast background of prematurely terminated RNAs destined for decay^12^. Precursor (pre-) spliceosomal U rich small nuclear RNAs (snRNAs, hereafter), U1, U11, U2, U12, U4 and U5 are transcribed by Pol II^16^. Following a first round of co-transcriptional processing, including 5’ end capping and 3’ end processing by the Integrator complex, snRNAs are exported to the cytoplasm after temporarily transiting through Cajal bodies, where a specific snRNA export complex is assembled^17^. While nuclear export of mRNA relies on export factor NXF1–NXT1 brought to the processed transcripts via the TREX and TREX-2 complexes^18,19^, nuclear export of snRNAs requires the RNA effector PHAX (Fig. 1a), which connects the CBC-bound transcript with the nuclear export factor CRM1 (also known as XPO1), itself bound to the small GTPase Ran and GTP (RanGTP)^20^. The snRNA export complex therefore comprises the CBC, PHAX, CRM1 and RanGTP but its assembly may require additional factors^21,22^. Mechanistically, RanGTP-bound CRM1 recruits PHAX by binding to its leucine-rich nuclear export sequence (NES) (Fig. 1a,b)^20,23^. The snRNA export complex then translocates through the nuclear pore complex (NPC) to the cytoplasm where it disassembles due to GTP hydrolysis triggered by Ran activating factors (Fig. 1b)^20,23^. Control of this process involves the nuclear phosphorylation of the phosphorylation site cluster 2 (ST2) of PHAX (Fig. 1a) by casein kinase 2 (CK2), which is essential for stable complex formation and export^20,24^. In the cytoplasm, snRNA export complex disassembly is further promoted by dephosphorylation of PHAX by protein phosphatase 2A (PP2A)^20,24^.

**Figure 1.**
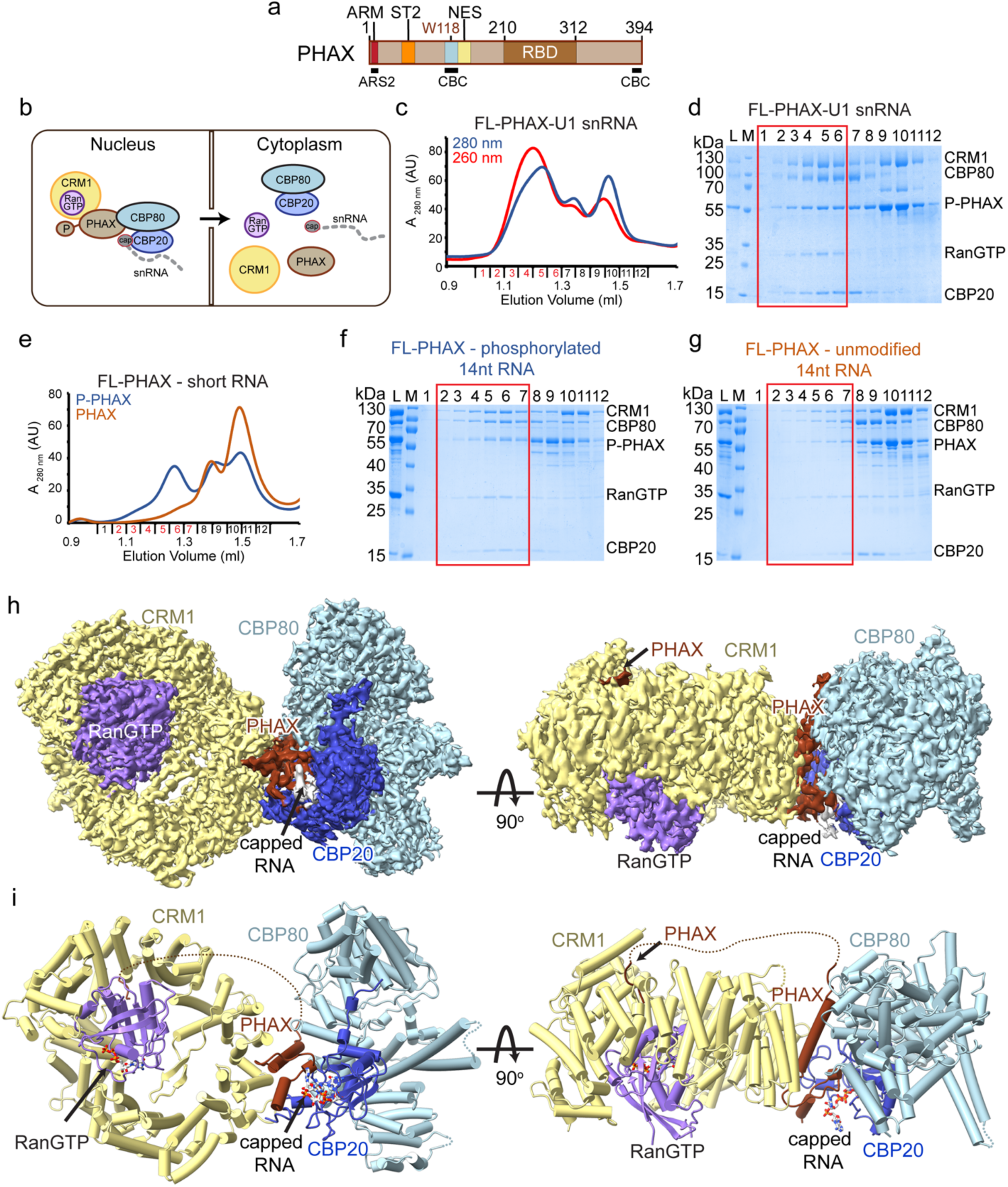
Structural characterization of the snRNA export complex **a.** Schematic representation of the PHAX domain structure. ARM: ARS2 recognition motif, ST2: phosphorylation site cluster 2, NES: nuclear export signal, RBD: RNA binding domain. The positions of the ARS2 (ARM) and CBC binding regions (W118) are shown. **b.** A schematic model of snRNA export mediated by the snRNA export complex assembled in the nucleus and disassembled in the cytoplasm. **c,d.** Superdex 200 gel filtration elution profile and SDS-PAGE analysis of fraction 1-12 of the snRNA complex reconstituted with capped U1 snRNA. The five proteins co-elute in the first peak highlighted with the red rectangle in **d**. L indicates input sample loaded onto the column. M: Mw marker **e.** Overlay of Superdex 200 gel filtration elution profiles of the snRNA complex reconstitutions with a short capped 14nt RNA using phosphorylated (P-PHAX) or unmodified PHAX. The complex elutes in fractions highlighted in red. **f,g.** SDS-PAGE analysis of fractions 1-12 of the gel filtration elution profiles shown in **e**. The red rectangle shows fraction containing the snRNA export complex. In panel **g**, the complex formation is less efficient and CBC co-elutes mostly with PHAX in fractions 8 and 9. **h.** Cryo-EM map used to build the snRNA export complex structure containing a 14nt capped RNA. Map is coloured according to molecules location in the structure: CBP80 is in light blue, CBP20 in blue, capped RNA in grey, PHAX in dark brick, CRM1 in yellow and Ran in violet. **i.** Ribbon representation of the snRNA export complex. RNA and GTP are shown as sticks.

The specificity of PHAX for short RNAs is speculated to depend on its competition with the hnRNP C protein, which selectively binds as a tetramer to transcripts with relatively long (>200–300 nts) and unstructured 5’ proximal regions^25,26^. hnRNP C may in this way restrict PHAX binding to short RNAs and to the CBC^25,27^. Finally, the PHAX-CRM1 pathway has also been implicated in the nuclear export of a small class of m^7^G-capped pre-microRNAs^28^ as well as histone H2AX mRNA^29^.

The human PHAX protein consists of 394 amino acid residues and is predicted to be mostly intrinsically disordered, with the exception of its RNA binding domain (RBD) that binds single-stranded RNA without sequence specificity (Fig 1a)^30,31^. PHAX can form a CBCAP complex with CBCA^32^ and *in vitro*, the N-terminally located ARM motif of PHAX binds to the effector domain of ARS2^7,33^. However, in the presence of ARS2, PHAX is unable to directly bind to the CBC due to steric competition by ARS2^7^. This suggests that ARS2 functions as a gatekeeper, preventing RNA-bound CBC to commit to RNA export until appropriate conditions are met. The cryo-EM structure of the CBC-PHAX (CBCP) complex revealed that the W118-containing helix of PHAX, as well as its C-terminus, directly binds to the CBC^7^. Consistent with a possible competition between ARS2 and PHAX for CBC binding, the PHAX C-terminus binds in the groove between CBP20 and CBP80 that also accommodates the ARS2 C-terminus^7^. Exactly how this relates to RNA sorting is currently unclear.

Here, we reconstitute *in vitro* the core snRNA export complex comprising capped RNA, CBC, phosphorylated full-length PHAX and CRM1-RanGTP, an example of a post-fate decision committed complex, and we confirm that PHAX phosphorylation biochemically stabilizes the complex. The cryo-EM structure at 2.45 Å resolution shows how synergistic interactions allow PHAX to simultaneously bridge the CBC and CRM1 while reinforcing dinucleotide cap recognition and CRM1 binding. Unexpectedly, we also find that a conserved two-residue motif of PHAX binds to a distant region of CRM1, suggesting that the adjacent phosphorylated ST2 cluster of PHAX could interact with a prominent basic patch of RanGTP. These observations are supported by mutagenesis of key PHAX residues *in vitro* and in cells. The Ran basic patch was previously shown to be important for binding of the RNA component of export complexes for tRNA, pre-miRNA and HIV RRE with their respective export adaptors EXPO-T, EXPO-5 and CRM1^34–36^. Thus, PHAX phosphorylation may not only increase export complex stability but also enable favourable competition with other CRM1 RNA cargoes, in a potentially regulatable fashion^20^.

## Results

### Reconstitution and cryo-EM structure of the snRNA export complex

To understand the molecular details underlying the role of PHAX within the snRNA export complex, we sought to determine its structure using single-particle cryo-EM. The snRNA export complex was reconstituted with purified CBC, *in vitro* CK2-phosphorylated full-length PHAX, exportin CRM1 together with the catalytic mutant of its co-factor Ran (Q69L) bound to GTP and an *in vitro* transcribed and capped U1 snRNA (164 nts). Phosphorylation of PHAX by CK2 was verified by gel shift assays and mass-spectrometry revealing up to 5 phosphorylation sites (4 within ST2) (Supplementary Fig. 1a,b). The complex was purified using size exclusion chromatography where all the components co-eluted in a peak with the expected apparent stoichiometry (Fig. 1c,d). An equivalent complex could also be reconstituted with a short capped-RNA containing only 14 nucleotides (m^7^GpppAAUCUAUAAUAGCA), suggesting that specific snRNA features might not be required for complex formation *in vitro* (Fig. 1e,f, Supplementary Fig. 2a,b). As expected, when PHAX was not phosphorylated, the snRNA export complex formed with significantly lower efficiency (Fig. 1e-g). A preliminary cryo-EM characterization of the two complexes, containing either the U1 snRNA or the short RNA, resulted in essentially identical reconstructions, where the bulk of the RNA was not visible. Here we describe the structure obtained using the sample containing the short capped-RNA. The structure of the complete complex was obtained at an overall resolution of 2.8 Å, limited by slight flexibility between the CRM1-RanGTP and CBC-PHAX moieties. Focussed refinement on each moiety separately improved the resolution to around 2.45 Å (Fig. 1h,i, Supplementary Fig. 2, Supplementary Table 1).

The cryo-EM map covers all the snRNA export complex components (Fig. 1h,i) with CBC, formed of subunits CBP80 and CBP20, and the export factor, consisting of CRM1 and RanGTP, adopting essentially the same structures as reported previously^8,37^. These two lobes of the structure are connected by PHAX. In our previous structure of the CBC-PHAX complex bound to a 12 nt capped RNA, only residues 114-132 of PHAX were visible, forming a short tryptophan-containing helix directly interacting with CBP80^7^ (Fig. 1a). Now, in the presence of CRM1-RanGTP, a larger region of PHAX is folded, encompassing residues 112-162. The previously observed W118-containing PHAX helix is extended to include the NES, which binds to CRM1, thus rigidly bridging the RNA-binding CBC to the export factor. The downstream segment of PHAX (residues 140-162) is also structured, forming contacts with both CBP20 and the RNA cap structure, while the architecture of the complex also enables direct contacts between CRM1 and CBP20. Finally, a second region of PHAX, spanning N-terminal residues 55-60, is observed to directly bind between HEAT repeats 5 and 6 of CRM1. Concerning the RNA, only 3 nucleotides of the cap structure are well-defined, indicating flexibility of the rest of the RNA molecule and the RBD domain of PHAX is also not visible in the map. The structure thus reveals the key role of PHAX^112–162^ in recruiting CRM1-RanGTP to the CBC-bound capped RNA.

### PHAX mediates the interaction between the CBC and CRM1

The largely conserved PHAX residues 116-136 form a long helix that interacts with both the CBC and CRM1 (Fig. 2a-c). The C-terminal half of the helix includes the NES and its residues dock into corresponding hydrophobic pockets of the NES-binding cleft of CRM1 between HEAT repeats 11 and 12 (Fig. 2c,d). While V130, L134, L137 and M139 bind CRM1 in a canonical way, the first residue of the PHAX NES sequence (Φ0) is a conserved glutamine (Q127) rather than a hydrophobic residue. This may weaken the PHAX NES-CRM1 interaction as compared to a canonical NES^38^ (Fig. 2b). CRM1 binding also includes hydrogen bonds formed between PHAX residues D128 and L137 and CRM1 residues H558 and K568 (Fig. 2d).

**Figure 2.**
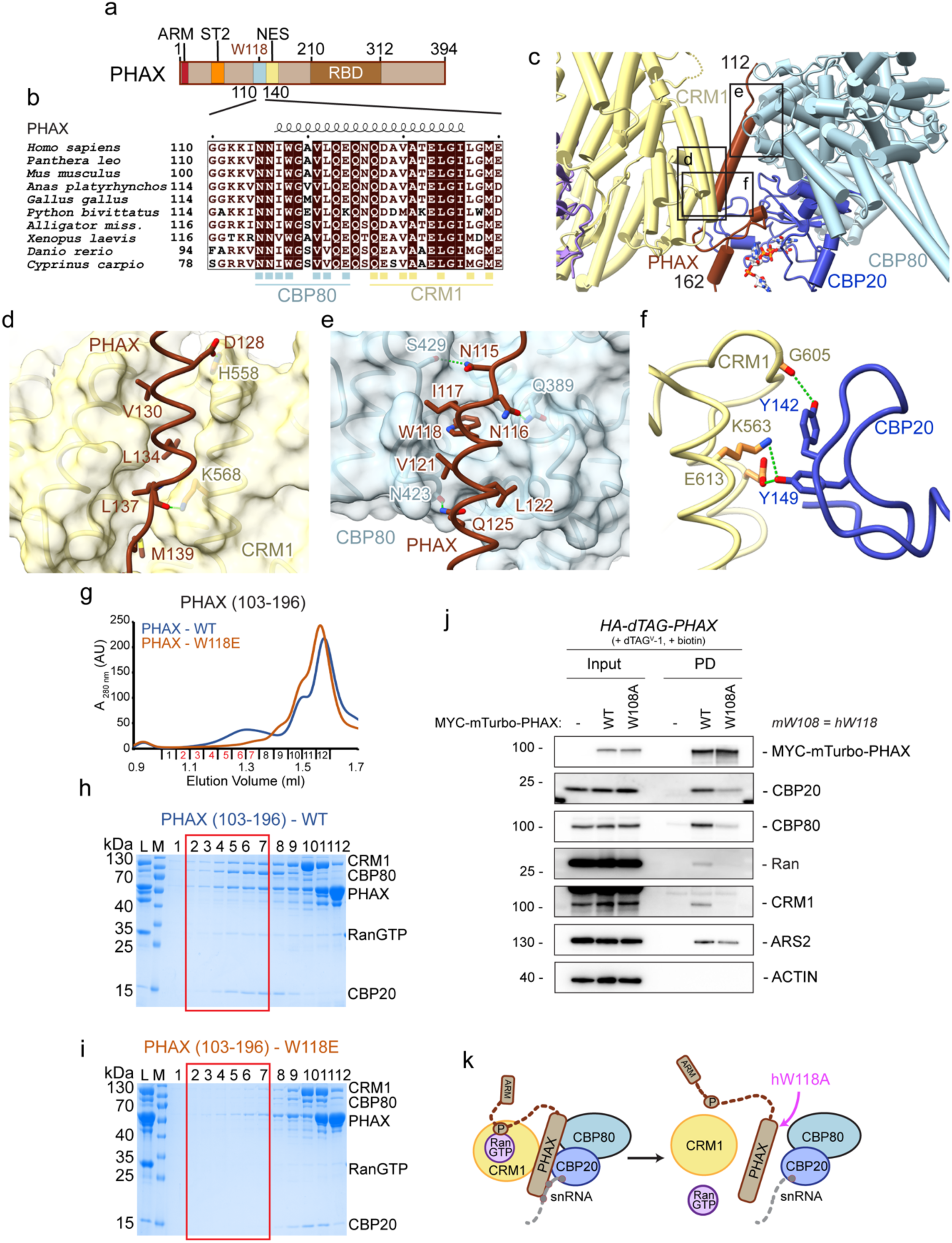
PHAX-mediated contacts between CBC and CRM-1 **a.** Schematic representation of the PHAX domain structure. ARM: ARS2 recognition motif, ST2: phosphorylation site cluster 2, NES: nuclear export signal, RBD: RNA binding domain. **b.** Sequence alignment of PHAX proteins covering the CBC-binding and NES regions. Identical residues are in brown boxes. Blue squares indicate residues involved in the interaction with CBC and yellow squares show residues interacting with CRM1 **c.** Ribbon representation the CBC-CRM1 interactions mediated by PHAX and CBP20. The three black rectangles position the close-up views shown in panels **d-f**. **d.** Details of the interactions between the PHAX helix (residues 117-136, dark brick) and CRM1 shown as surface (yellow). The NES residues of PHAX insert into hydrophobic pockets on CRM1. Several charged contacts with CRM1 are shown. **e.** Details of the hydrophobic and charged interactions between the PHAX helix (residues 117- 136, dark brick) and CBP80 shown as surface (light blue). **f.** Molecular details of the interactions between the CRM1 (yellow) and CBP20 (dark blue). **g.** Overlay of Superdex 200 gel filtration elution profiles of the snRNA complex reconstitutions with using either WT or W118E mutant of MBP-PHAX^103-196^ and a short capped 14nt RNA. The WT complex elutes in fractions highlighted in red. **h,i.** SDS-PAGE analysis of fractions 1-12 of the gel filtration elution profiles shown in **g**. The red rectangle shows fraction containing the snRNA export complex. In panel **i**, MBP-PHAX^103-196^ W118E is unable to form the complex with CBC and CRM1-RanGTP. L indicates input sample loaded onto the column. M: Mw marker **j.** Western blotting analysis of streptavidin pulldowns from lysates of *HA-dTAG-PHAX* cells stably expressing MYC-mTurbo tagged PHAX^WT^ or PHAX^W108A^ with the parental cell line (-) serving as a negative control. Cells were treated with dTAG^V^-1 for 4 hours to deplete endogenous HA-dTAG-PHAX protein and treated with biotin for a further 4 hours to induce proximity labelling. Input (left) and streptavidin-enriched pulldown (PD, right) samples were probed with antibodies against MYC, CBP20, CBP80, Ran, CRM1 and ARS2 with ACTIN serving as an input loading control **k.** A schematic model of the impact of hW118A mutation on the snRNA export complex assembly.

The N-terminal half of the same PHAX helix binds to the CBC as also reported in our previous CBC-PHAX-RNA structure^7^, with the W118 residue inserting into the conserved pocket formed by CBP80 residues A392, H420, F424 and W428 (Fig. 2c,e). However, when PHAX is engaged within the snRNA export complex, several extra contacts occur between the two proteins, including hydrogen bonds formed between N115, N116 and Q125 of PHAX and S429, Q389 and N423 of CBP80 (Fig. 2e). Whereas the extreme C-terminus of PHAX (residues 390-394) was observed to bind in the groove between CBP80-CBP20 in the binary CBC- PHAX complex^7^, it is not visible in the snRNA complex structure, the groove being empty. In the snRNA export complex, the CBC and CRM1 also directly interact with each other via several hydrogen bonds between Y142 and Y149 of CBP20 and K563, G605 and E613 of HEAT repeats 12 and 13 of CRM1 (Fig. 2f).

PHAX, NELF-E, NCBP3 and ZC3H18 all recognise the same surface on the CBC via their conserved tryptophan-containing helices^7^. Furthermore, the mRNA export factor ALYREF interacts with a neighbouring CBC surface^39^. Comparison of our snRNA export complex structure with that of the CBC bound to the RRM domain of ALYREF, revealed that ALYREF binding to CBC is incompatible with the CBC binding to PHAX and CRM1 (Supplementary Fig. 3a-c). This mutual exclusivity may thus contribute to directing snRNAs and mRNAs into distinct pathways for nuclear export.

### CBC binding by the tryptophan-containing helix of PHAX is critical for snRNA complex assembly

Since the PHAX core region observed in the cryo-EM structure includes residues 112- 162, we tested whether a corresponding PHAX fragment would be sufficient to form the snRNA export complex. Indeed, PHAX^103–196^ lacking the upstream CRM1-binding sequence (residues 55-60, see below), the ST2 and the RBD still formed the complex, albeit with lower CRM1 stoichiometry as compared to full-length phosphorylated PHAX (Fig. 2g,h). We previously showed that the PHAX W118E mutation was sufficient to disrupt PHAX^103–196^ binding to the CBC^7^. Interestingly, W118E mutation also impeded snRNA export complex formation (Fig. 2g,i), showing that the interaction between the W118-containing helix and the CBC remains critical within the complete export complex.

To test the importance of the PHAX W118-containing helix in the formation of the snRNA export complex *in vivo*, we utilised a proximity labelling pulldown approach to explore the recruitment of proteins to wildtype (WT) or mutated PHAX. We first tagged endogenous PHAX alleles in mouse embryonic stem (mES) cells with a 2xHA-FKBP-V (HA-dTAG) degron tag to allow for the rapid depletion of HA-dTAG-PHAX proteins upon the addition of the dTAG^V^-1 ligand. This was efficiently achieved following 2 hours of dTAG^V^-1 treatment, with no consequential effects on steady state levels of CBCA or CRM1-Ran proteins (Supplementary Fig. 3d). We then utilized the PiggyBac transposase system^40^ to stably introduce MYC-mini-Turbo (mTurbo) tagged PHAX^WT^ or PHAX^W108A^ (mouse equivalent of W118) into the HA-dTAG-PHAX cells. The mTurbo proximity labelling system allows for *in vivo* biotinylation of proteins in the vicinity (1-10 nm) of the fusion protein upon the addition of biotin to media. Following depletion of the endogenous HA-dTAG-PHAX protein and subsequent addition of biotin, MYC-mTurbo-PHAX^WT^ and PHAX^W108A^ efficiently biotinylated proteins when compared to cells lacking mTurbo (Supplementary Fig. 3e). Having validated the system, we proceeded to purify *in vivo* biotinylated proteins from cells expressing MYC-mTurbo-PHAX^WT^ or PHAX^W108A^ by streptavidin-enrichment.

Western blotting analysis revealed that PHAX^WT^ pulldowns were enriched for CBC (CBP80, CBP20), ARS2, CRM1 and Ran proteins as compared to an untagged control (Fig. 2j). However, the PHAX^W108A^ mutant showed a marked decrease in CBC proteins and a complete loss of CRM1 and Ran (Fig. 2j,k). We speculate that the residual association of CBC with PHAX^W108A^ might be mediated by ARS2, bridging the ARM motif of PHAX and the groove between CBP20 and CBP80^7,9^, which instead, is insufficient for the recruitment of CRM1-RanGTP. Such involvement of ARS2 in the complex will be discussed later. For now we conclude that the W118-mediated interaction between PHAX and CBC is crucial for efficient formation of the snRNA export complex (Fig. 2k).

### PHAX is intimately involved in RNA cap recognition

The snRNA export complex structure reveal that conserved PHAX residues 140-162, downstream of the PHAX^116–136^ bridging helix, engage in specific contacts with the RNA cap structure, CBP20 and CRM1 (Fig. 3a-c). Details of m^7^GTP cap-moiety recognition by CBP20, in the context of CBC alone, are well-established^8,10^ (Supplementary Fig. 4a) and in the presence of PHAX, the methylated guanosine and the α-phosphate of the cap are bound by CBP20 as previously reported. However, in addition, we found that the conserved E150 residue of PHAX inserts into the cap-binding site of CBP20, forming hydrogen bond interactions with R112 and Y43 residues of CBP20. Since these two residues are key for the specific interaction with m^7^G, PHAX likely strengthens cap recognition by CBP20 (Fig. 3d).

**Figure 3.**
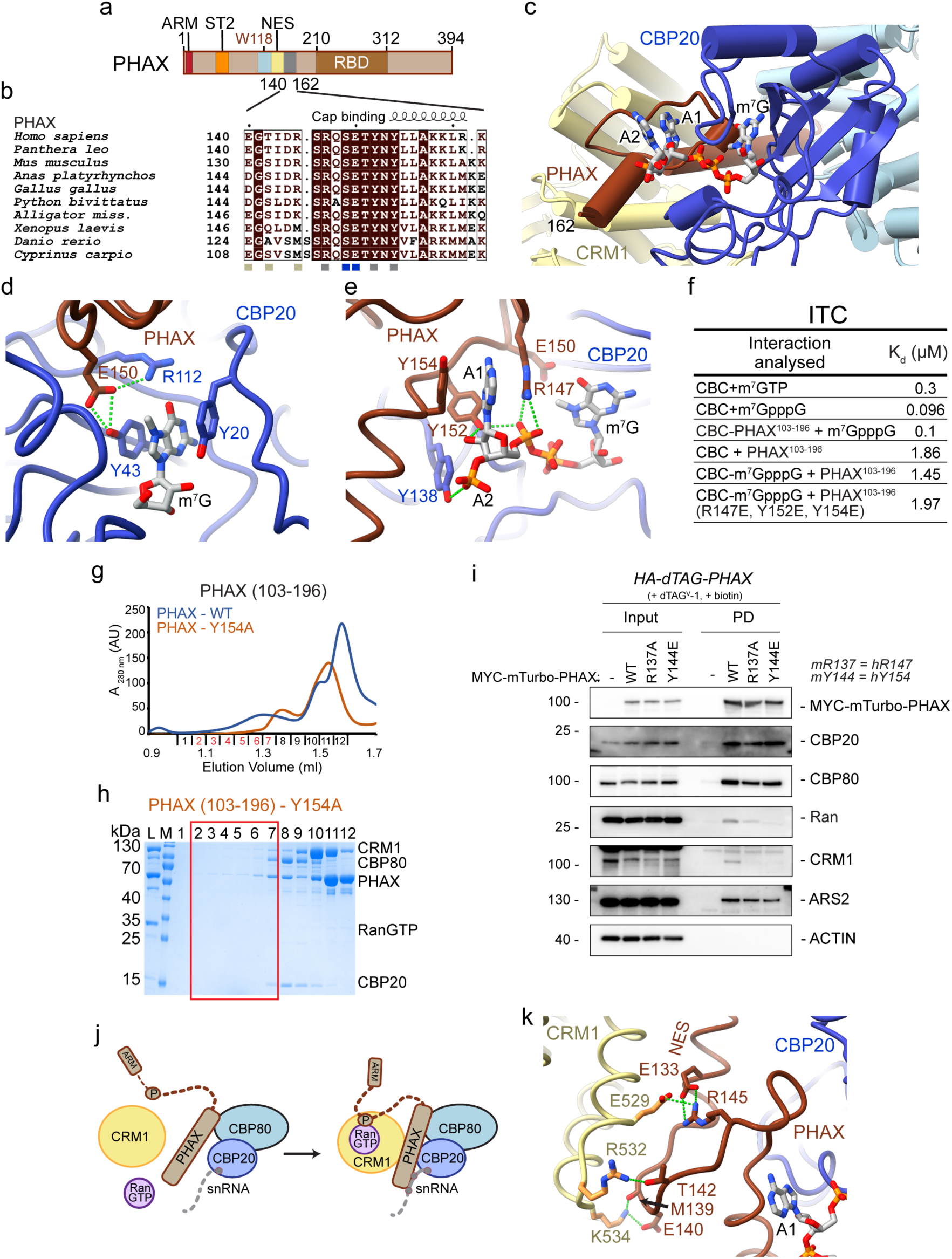
RNA recognition by PHAX **a.** Schematic representation of the PHAX domain structure. ARS2 recognition motif, ST2: phosphorylation site cluster 2, NES: nuclear export signal, RBD: RNA binding domain. **b.** Sequence alignment of PHAX proteins covering the cap- and CRM1-binding regions. Identical residues are in brown boxes. Blue squares indicate residues involved in the interaction with cap and yellow squares show residues interacting with CRM1 **c.** Ribbon representation the RNA recognition by the snRNA export complex. **d.** Details of the interactions of PHAX E150 with key CBP20 cap-binding residues. **e.** Details of the specific recognition of the first-transcribed residues by PHAX. **f.** Summary of the ITC measurements of binding affinities between CBC, PHAX^103-196^ and the cap structure **g.** Overlay of Superdex 200 gel filtration elution profiles of the snRNA complex reconstitutions with using either WT or Y154A mutant of MBP-PHAX^103-196^ and a short capped 14nt RNA. The WT complex elutes in fractions highlighted in red. **h.** SDS-PAGE analysis of fractions 1-12 of the gel filtration elution profiles shown in **g.** MBP- PHAX^103-196^ Y154A is unable to form the complex with CBC and CRM1-RanGTP. The fractions corresponding to the WT complex elution peak are highlighted with red rectangle. L indicates input sample loaded onto the column. M: Mw marker **i.** Western blotting analysis of streptavidin pulldowns from lysates of HA-dTAG-PHAX cells stably expressing MYC-mTurbo tagged PHAX^WT^, PHAX^R137A^ or PHAX^Y144E^ as described in Fig 2j. **j.** A schematic model of snRNA export formation based on RNA- and CRM1-RanGTP- triggered folding of PHAX. **k.** Details of direct interactions between CRM1 and the PHAX region downstream NES.

A key difference from cap binding by CBC alone, is the strong additional binding of PHAX to the first-transcribed RNA nucleotide. In the CBC-cap analog (m^7^GpppG) complex structure, determined in the absence of PHAX^8^, the first-transcribed nucleotide (here G1) stacks with the Y138 residue of CBP20 (Supplementary Fig. 4a). Moreover, studies employing the Y138A mutant showed that this residue confers additional binding affinity for dinucleotide cap analogues with a purine as first-transcribed nucleotide^41^. However, in the presence of PHAX, the first-transcribed nucleotide (here A1) is shifted away from CBP20 and forms multiple interactions with PHAX (Fig. 3e, Supplementary Fig. 4a). Importantly, A1 is sandwiched between PHAX residues R147 and Y154. In addition, R147 together with Y152 interacts with the phosphate Pγ and the ribose moiety of A1 (Fig. 3e). This results in a slight movement of Pβ and Pγ, and loss of a hydrogen bond between Pβ and CBP20 R127 (Supplementary Fig. 4a). Y138 of CBP20 now binds the phosphate group of the following nucleotide (Fig. 3e, Supplementary Fig. 4a). This CBC-PHAX cap recognition is likely compatible with a Cap-1 structure bearing an additional 2′O ribose methylation of A1 and maybe reinforced by eventual m6 methylation of A1.

### Cap recognition by PHAX requires the presence of CRM1-RanGTP

The snRNA export complex structure reveals that in the presence of CRM1 and RanGTP, PHAX contributes significantly to RNA cap binding (Fig. 3c-e). However, in our previous cryo-EM structure of full-length PHAX bound to the CBC and a 12 nt capped RNA, no contacts of PHAX with the cap were visible^7^. These observations suggest that PHAX recognition of RNA might only occur when residues 140-162 are stabilised by the presence of CRM1-RanGTP. To address this issue, we measured the binding affinities between CBC, PHAX^103–196^ and the cap structure by isothermal calorimetry (ITC). In doing so, we found that CBC binds m^7^GTP with a *K*_d_ of 300 nM (Fig. 3f, Supplementary Fig. 4b). For the cap analogue m^7^GpppG, the measured *K*_d_ was 96 nM (Fig. 3f, Supplementary Fig. 4c). This higher affinity for the cap analog is consistent with previous observations that the first-transcribed G stacks with Y138 of CBP20^8,10^. Importantly, when CBC was in complex with PHAX^103–196^, that bears residues sufficient for CBC and cap binding, the affinity for the cap analogue remained unchanged, suggesting that PHAX does not contribute to RNA binding in this context, as observed structurally (Fig. 3f, Supplementary Fig. 4d). Similarly, the presence of the second nucleotide of the cap structure did not result in a significant increase of the affinity of CBC to PHAX^103–196^ and the triple mutation of the three key A1-binding residues of PHAX (R147E, Y152E, Y154E) had essentially no impact on PHAX binding to the CBC in presence of the cap analogue (Fig. 3f, Supplementary Fig. 4e-g). Thus, in absence of CRM1-RanGTP, the addition of PHAX to the CBC does not result in a measurable binding affinity change to the cap structure, suggesting that the corresponding regions of PHAX are disengaged.

### Cap recognition by PHAX is required for the recruitment of CRM1-RanGTP

Next, we investigated the role of the capped RNA in snRNA export complex formation. While CRM1/RanGTP-triggered folding of PHAX^140-162^ is needed for the enhanced recognition of capped RNA, reciprocally, the structure also indicated that cap-dependent folding of PHAX^140-162^ is likely required to achieve an efficient interaction between CBC-PHAX and CRM1-RanGTP. Indeed, using size exclusion chromatography and mass photometry, we could show that snRNA export complex formation depends on the presence of capped RNA, with at least a dinucleotide cap being required (Supplementary Fig. 5a-c). Furthermore, triple mutation, R147E, Y152E, Y154E, within the cap-binding region of PHAX^103-196^ disrupted the complex (Supplementary Fig. 5d,e). The same negative impact was observed even for the single Y154A mutation (Fig. 3g,h). Our parallel *in vivo* analysis, employing PHAX R137A and Y144E mutations (mouse equivalents of R147A or Y154E), showed significantly reduced proximity ligation of PHAX with CRM1 and Ran, while labelling of CBC and ARS2 remained unchanged (Fig. 3i). We therefore conclude that the strong interaction of the first transcribed nucleotide with PHAX is required for efficient recruitment of CRM1 and RanGTP to the complex.

snRNA export complex formation thus requires a synergistic, mutually dependent assembly of all its components. It has been established that efficient NES recognition by CRM1 requires the presence of Ran-GTP^42^ and that efficient cap binding by CBP20 depends on CBP80^8^. PHAX is key for bridging these two modules and its proper conformation depends on the presence of CRM1-RanGTP, CBC and capped RNA (Fig. 3j). An important role in the structuring of PHAX is probably also played by residue R145, which connects the PHAX region bound to RNA and CBP20 to the PHAX helix and CRM1, through charged interactions with PHAX E133 and CRM1 E529 (Fig. 3k). PHAX residues M139, E140 and T142 also form additional polar contacts with CRM1 residues R532 and K534, thus strengthening the NES- CRM1 interface.

### PHAX phosphorylation enhances stability and specificity of the snRNA export complex

PHAX phosphorylation mediated by CK2 is required for the stable formation of the snRNA export complex^20,24^. PHAX contains 8 putative phosphorylation clusters (ST) but only the second cluster (ST2, residues 64-76) (Fig. 4a,b) is essential for snRNA export complex assembly and export^24^. Conducting gel filtration experiments with unmodified PHAX, we confirmed that the snRNA export complex forms with significantly lower efficiency (Fig. 1e- g). Similarly, we found that PHAX^103-196^, that lacks ST2 and thus is not phosphorylated but encompasses the part of PHAX at the CBC-CRM1 interface, can still form the snRNA export complex, albeit less efficiently (Fig. 2g-i). Equivalent results were obtained using mass photometry (Supplementary Fig. 6a). How phosphorylation of ST2 enhances complex assembly and promotes export is therefore an important question.

**Figure 4.**
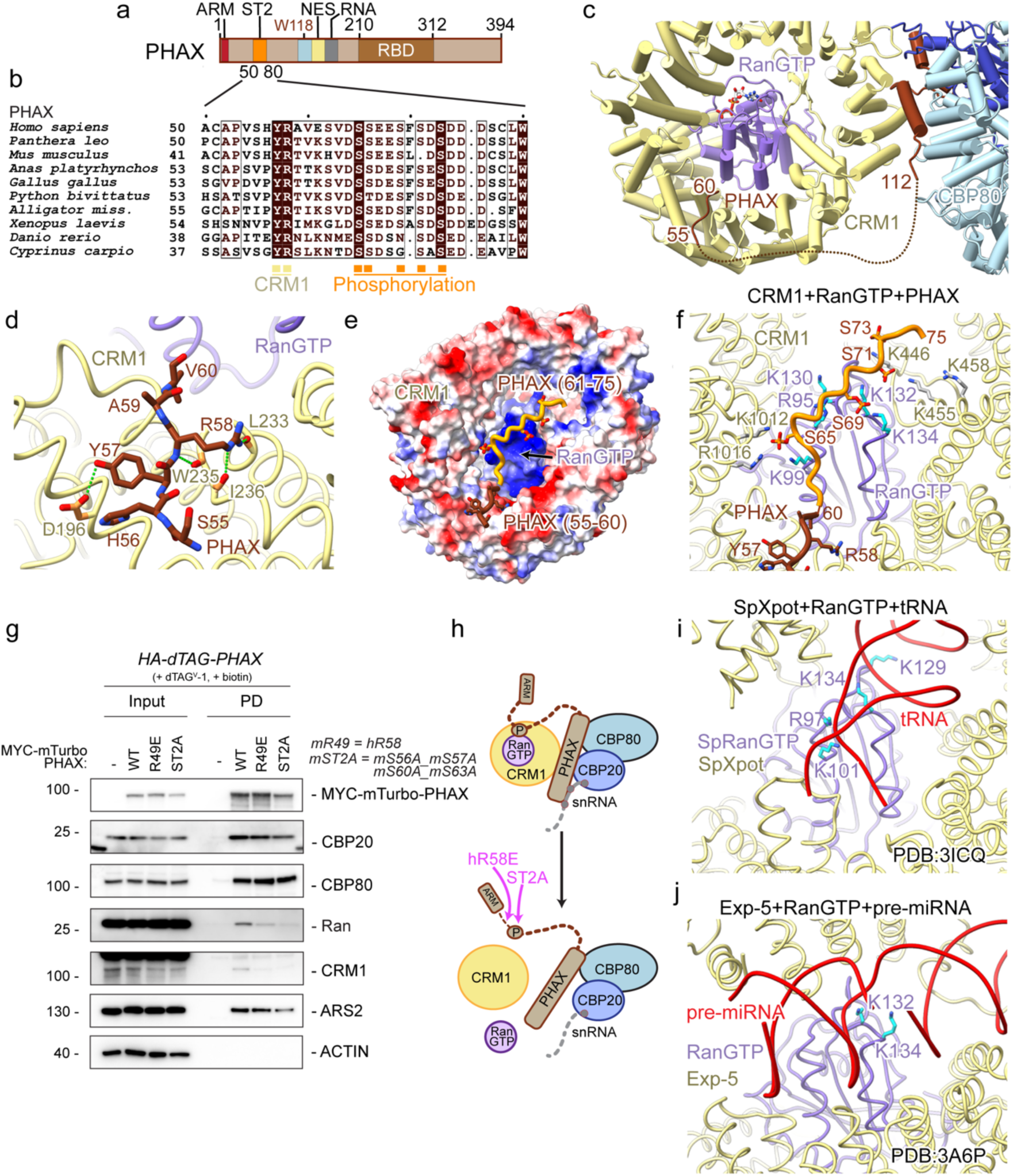
PHAX N terminus connects CRM1-RanGTP **a.** Schematic representation of the PHAX domain structure. ARM: ARS2 recognition motif, ST2: phosphorylation site cluster 2, NES: nuclear export signal, RBD: RNA binding domain. **b.** Sequence alignment of PHAX proteins covering the CRM1-binding and phosphorylation regions. Identical residues are in brown boxes. Orange squares indicate potential phosphorylated residues and yellow squares show residues interacting with CRM1 **c.** Ribbon representation of the interaction between CRM1 and the N terminus of PHAX. **d.** Details of the interactions between PHAX^56-60^ and CRM1 showing charged contacts between the two proteins. **e.** Electrostatic surface representation of the CRM1-RanGTP. PHAX^55-60^ is shown in brown as sticks and the AlphaFold3 model of following the ST2 region is in orange with phosphorylated serines as sticks binding over the basic surface of Ran. **f.** Cartoon representation of the CRM1-RanGTP-PHAX complex. PHAX^55-60^ is shown in brown as sticks. The AlphaFold3 model of following the ST2 region is shown in orange with phosphorylated serines as sticks positioned in proximity of positively charged residues of the basic surface of Ran. **g.** Western blotting analysis of streptavidin pulldowns from lysates of HA-dTAG-PHAX cells stably expressing MYC-mTurbo tagged PHAX^WT^, PHAX^R49E^ or PHAX^ST2A^ as described in Fig 2j. **h.** A schematic model of the impact of hR58E and ST2A mutation on the snRNA export complex assembly. **i.** Contacts between the basic surface of *S. pombe* RanGTP and tRNA within the tRNA nuclear export complex^34^. **j.** Contacts between the basic surface of RanGTP and pre-miRNA within the pre-miRNA nuclear export complex^35^.

Even though PHAX^112-162^ is the key region bridging CBC-RNA to CRM1-RanGTP, the cryo-EM map also exhibited extra density between CRM1 HEAT repeats 5 and 6. Interestingly, AlphaFold2^43^ predictions of the CRM1-RanGTP-PHAX complex, show, with high confidence, binding of PHAX^56-60^ to exactly this position (Supplementary Fig. 6b-d). The predicted position of this peptide fits very well into the extra cryo-EM density (Supplementary Fig. 6e) and two confidently predicted and conserved PHAX residues, Y57 and R58 (Fig. 4b), engage in several charged contacts with CRM1 residues (Fig. 4c,d). Intriguingly, PHAX^56-60^ is immediately upstream of the ST2 phosphorylation cluster, which also includes a number of acidic residues (64-DSSEESFSDSDDD) (Fig. 4a,b), typical of a multiple CK2 phosphorylation site^44,45^. AlphaFold3 (AF3)^46^ is capable of making predictions using posttranslational modifications, such as phosphorylation. Whilst AF3 does not make a unique, confident prediction of the position of the ST2 pSer residues, all models displayed the ST2 peptide (which is pinned on one side by the Y57-R58 binding) lying on a strongly basic surface of CRM1-RanGTP (Fig. 4e,f, Supplementary Fig. 6f). Bearing in mind that ST2 phosphorylation may be heterogeneous, this would be consistent with a possibly ‘fuzzy’, electrostatically-driven interaction between ST2 and this region of CRM-RanGTP. As an indication of how the pSer (and other acidic residues in ST2 of PHAX) could interact to the basic surface formed by both CRM1 and Ran, pSer65 is spatially close to CRM1 residues K1012, R1016 and Ran residue K99. Moreover, pSer69 and 71 are close to Ran K132, K134. Additionally, Ran R95, K130 and CRM1 K446, K455, R458 would be in the vicinity of the ST2 peptide (Fig. 4f). Indeed, further examination of the CryoEM map showed weak extra density to support this suggestion (Supplementary Fig. 6g).

The importance for snRNA export complex formation of both PHAX^55-60^ binding to CRM1 and ST2 phosphorylation was tested *in vivo*, employing mutation of equivalent residues in the mouse PHAX protein. These mutants (mR49E and mS56A-S57A-S60A-S63A) significantly reduced recruitment of CRM1 and RanGTP to PHAX, while having little impact on CBCA association (Fig. 4g,h). We therefore conclude, that the interaction between PHAX^55-60^ and CRM1, that likely enables proper positioning of the phosphorylated ST2 region on the basic surface on RanGTP, is required for efficient CRM1-RanGTP recruitment to the complex. Within tRNA-, pre-miRNA- and HIV RRE- export complexes, the basic surface of Ran forms direct contacts with RNA (Fig. 4i,j)^34–36^. It is thus possible that in addition to enhancing export complex stability, PHAX phosphorylation also enables favourable competition with noncognate CRM1 RNA cargos.

### ARS2 is tenuously associated with the snRNA export complex via the PHAX ARM motif

PHAX can form a ternary complex with CBCA^9,32^. Within this complex, the ARM of PHAX (9-EDGQL-13) binds the effector domain of ARS2^7^ (Fig. 5a) likely in a manner similar to that observed in the crystal structure of ARS2 bound to the ARM motif of the *S. pombe* Red1 protein^13^. Reciprocally, the ARS2-PHAX interaction is lost upon mutation or deletion of the effector domain of ARS2^33,47^. In ternary complexes, ARS2 shields direct binding of PHAX, as well as the effectors NCBP3 and ZC3H18, to the CBC by sterically excluding them from the CBP20-CBP80 groove and the tryptophan helix binding sites^7^. Based on these data, the relationship of ARS2 with the snRNA export complex is unclear and we therefore used size exclusion chromatography with purified proteins to assess whether ARS2 could be a component of the export complex.

**Figure 5.**
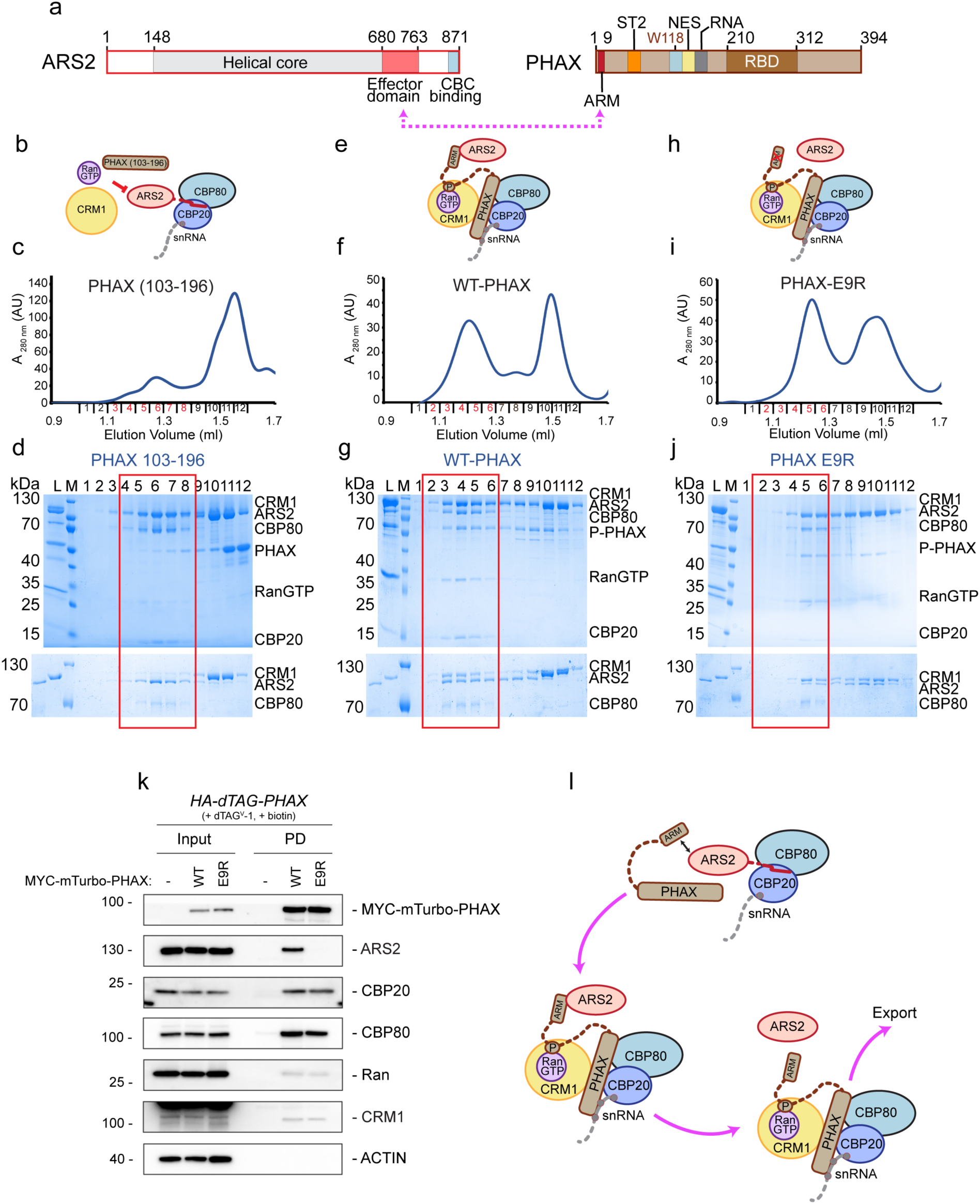
ARS2 interaction with the snRNA export complex. **a.** Schematic representation of the ARS2 and PHAX domain structure. The ARS2 effector domain recognises the ARM motif in PHAX N terminus. ARM: ARS2 recognition motif, ST2: phosphorylation site cluster 2, NES: nuclear export signal, RBD: RNA binding domain. **b.** Schematic representation of the interaction of the CBC-ARS2^147-871^-RNA complex with CRM1, RanGTP and PHAX^103-196^. **c.** Superdex 200 gel filtration elution profile of the mixture of the CBC-ARS2^147-871^-RNA complex with CRM1, RanGTP and PHAX^103-196^. The CBC-ARS2^147-871^-RNA complex elutes in fractions highlighted in red. **d.** SDS-PAGE analysis of fraction 1-12 of the gel filtration elution profile shown in **c**. Lower panel shows a 7% SDS-PAGE analysis of the same fractions enabling better resolution of CRM1 and ARS2^147-871^. The red square indicates fractions containing the CBC-ARS2^147-871^- RNA complex. CRM1, PHAX and RanGTP elute in later fractions. L indicates input sample loaded onto the column. M: Mw marker **e.** Schematic representation of the interaction of the CBC-ARS2^147-871^-RNA complex with CRM1, RanGTP and FL PHAX. ARS2 binds to PHAX via its ARM motif. **f.** Superdex 200 gel filtration elution profile of the mixture of the CBC-ARS2^147-871^-RNA complex with CRM1, RanGTP and FL PHAX. The snRNA export complex bound to ARS2^147-^ elutes in fractions highlighted in red. **g.** SDS-PAGE analysis of fraction 1-12 of the gel filtration elution profile shown in **f**. The red square indicates fractions containing the snRNA export complex bound to ARS2^147-871^. **h.** Schematic representation of the interaction of the CBC-ARS2^147-871^-RNA complex with CRM1, RanGTP and E9R mutant of FL PHAX. ARS2 binding to PHAX via its ARM motif is prevented. **i.** Superdex 200 gel filtration elution profile of the mixture of the CBC-ARS2^147-871^-RNA complex with CRM1, RanGTP and FL PHAX E9R. **j.** SDS-PAGE analysis of fraction 1-12 of the gel filtration elution profile shown in **i**. The red square indicates fractions where the snRNA export complex bound to ARS2^147-871^ eluted in **g.** Here, ARS2 becomes sub-stoichiometric and the snRNA complex elutes later in fraction 4-6. **k.** Western blotting analysis of streptavidin pulldowns from lysates of HA-dTAG-PHAX cells stably expressing MYC-mTurbo tagged PHAX^WT^ and PHAX^E9R^ as described in Fig. 2j. **l.** Model for the role of ARS2 in the snRNA export complex assembly. ARS2 binds to the groove formed at the CBP20-CBP80 interface. While it may interact with the ARM motif of PHAX it prevents its binding to CBC. In the presence of CRM1-RanGTP and capped RNA, PHAX displaces ARS2 from CBC. ARS2 remains bound to the complex via the ARM motif but might as well leave and associate with other CBC-effectors complexes.

First, we assembled and purified CBC, ARS2^147-871^ and capped RNA. This complex was incubated with CRM1, RanGTP and PHAX^103-196^, lacking the ARM motif but including sufficient information to form the snRNA complex (Figure 2g,h). In the presence of ARS2, the snRNA export complex could not be formed and the CBC remained bound only by ARS2. Due to the similar migration of CRM1 and ARS2^147-871^, this was shown by resolving the respective elution profiles on 7% SDS PAGE gels (Fig. 5b-d). Hence, when PHAX does not possess all its features, ARS2 blocks access to the CBC even in the presence of other snRNA export complex components. In contrast, when the CBC-ARS2^147-871^-RNA complex was incubated with CRM1, RanGTP and phosphorylated FL PHAX (which includes the ARM), the snRNA complex was formed efficiently, eluted earlier and contained ARS2^147-871^ (Fig. 5e-g). Thus, in this context, ARS2 no longer prevents PHAX from binding to the CBC. We considered that ARS2 could be connected to the complex by its effector domain binding to the ARM of PHAX or by its C-terminus binding into the CBP20-CBP80 groove^7,9^. To distinguish these possibilities, we used an FL PHAX variant with an E9R mutation within the ARM, abolishing binding to ARS2^147-8717^. When the CBC-ARS2^147-871^-RNA complex was incubated with CRM1, RanGTP and phosphorylated FL PHAX E9R, the snRNA complex was formed, but the presence of ARS2^147-871^ was significantly reduced (Fig. 5h-j). Thus, without a connection between ARS2 and PHAX, the former is excluded from the export complex, likely because the reinforced binding of PHAX to the CBC in this complex out-competes the CBC-binding of ARS2. In agreement, in a 3.85 Å resolution cryo-EM map of the snRNA export complex bound to ARS2, no density corresponding to ARS2 could be observed in the CBP20-CBP80 groove (data not shown).

To investigate a role of ARS2 in snRNA export complex formation *in vivo*, proximity labelling was performed with the PHAX E9R mutant, which revealed that the abrogated PHAX ARM-ARS2 interaction had no apparent impact on PHAX contacts with core snRNA complex subunits (Fig. 5k). This indicates that, in contrast to the NEXT complex subunit ZC3H18^7^, ARS2 is not essential for PHAX recruitment to CBC-bound RNA. While the exact role of the PHAX-ARS2 interaction remains to be elucidated, these data suggest that ARS2 may initially prevent PHAX from binding the CBC, but in the presence of CRM1-RanGTP and capped RNA, the PHAX-CBC complex can be stably formed. At this stage, ARS2 is excluded from the CBC, but may remain connected via the PHAX ARM (Fig. 5l).

## Discussion

For 25 years PHAX has been known to be crucial for snRNA export^20^, but the molecular details underlying this role have remained elusive. Here we report the cryo-EM structure of the PHAX-containing snRNA export complex, revealing several novel aspects of PHAX activity and shedding light on the competition between RNA effectors for CBC binding.

PHAX forms a complex with CBC-ARS2 named CBCAP^32^. We have recently characterised this complex and shown that PHAX binds directly to the CBC with both its W118-containing helix and its C-terminus, while also binding to ARS2 via its N-terminal ARM motif^7^. When PHAX is bound to CBC and capped RNA, only the N-terminal part of the W118- containing helix of PHAX (residues 114-132) is stably bound to CBP80^7^. However, in the presence of CRM1-RanGTP, the W118-containing helix is extended to include the PHAX NES, which binds to the canonical NES binding cleft of CRM1. Our *in vitro* and *in vivo* data demonstrate that the W118-mediated interaction of PHAX with CBC is required for snRNA export complex formation. Since CRM1 is crucial for mediating snRNA nuclear export, this interaction is likely indispensable. An equivalent CBC interaction with the W301-containing helix of ZC3H18 is similarly important for the function of the NEXT complex in RNA degradation^7^. The same surface on the CBC is also used by NCBP3 but the functional role of this interaction is not known^7^. The Trp-binding surface on the CBC is thus emerging as a key contributor to RNA fate determination through its interactions with competing effectors. In the future, it will be interesting to investigate whether other important CBC-associating RNA effectors, such as ZFC3H1 of the PAXT RNA decay connection, also use this CBC surface for their function.

In the structure of PHAX bound to CBC and capped RNA, no contacts of PHAX with RNA were observed^7^. Within the snRNA export complex, however, PHAX engages in strong contacts with the RNA cap structure, in particular, sandwiching the first transcribed nucleotide and reinforcing the binding of m^7^GTP by CBP20. The CBC-PHAX module binds tightly the entire cap structure through π-π or π-cation stacking interactions with the two cap nucleotides and numerous charged contacts that include also the triphosphate moiety. This cap recognition can likely accommodate 2′O ribose methylation (Cap-1 structure). Moreover, most snRNAs have an A as first transcribed nucleotide whose further methylation to m^6^Am could possibly enhance the stacking energy between PHAX residues R147 and Y154. However, this modification is reversible and whether it remains on the pre-snRNA is highly regulated as well as both cell-type and RNA specific^48^. While it was shown that the RNA binding domain of PHAX plays an important role in snRNA export^31^, this domain is not observed in the structure with a 14 nt RNA nor with U1 snRNA (data not shown). Instead, we show that efficient CRM1- RanGTP recruitment requires PHAX structuring, triggered by RNA, which is likely a mechanism to avoid unnecessary nuclear export of CBC and PHAX. Conversely, capped RNA is recognised by PHAX only in the presence of CRM1-RanGTP, which allows stronger binding of snRNA substrates once nuclear export is committed.

Efficient snRNA export complex formation requires PHAX phosphorylation by CK2^20,24^. PHAX contains eight putative phosphorylation clusters (ST), but only phosphorylation of the second cluster, ST2 (residues 64-76), is necessary for export complex formation^24^. Our mass spectrometry analysis of *in vitro* CK2-phosphorylated PHAX revealed five phosphorylated serines in the full-length protein and four in the PHAX^41-80^ peptide, which only contains ST2. This supports that ST2 is the primary CK2 phosphorylation site in PHAX, at least in *vitro*. In the snRNA export complex structure, in addition to its NES binding to CRM1, PHAX also interacts with a distant region of CRM1 via its residues 55-60, containing the conserved 57-YR motif. We propose that this contact positions a ‘fuzzy’, electrostatically driven interaction between the adjacent phosphorylated ST2 cluster and the prominent basic surface of RanGTP, which further strengthens CRM1-RanGTP recruitment into the complex. Phosphorylated ST2 may also shield the basic surface on Ran from nonspecific interaction with non-cognate RNA cargos. Indeed, within the snRNA export complex structure, the bulk of the RNA is not visible and PHAX recognises specifically the cap structure. In contrast, within the tRNA, pre-miRNA and HIV RRE export complexes, the cargo RNA molecules form direct contacts with the same basic surface of Ran^34–36^. It cannot be ruled out that some element of the snRNA also contacts this surface at some point.

Our structure shows that PHAX residues 112-162, the key region of for snRNA export complex assembly, only folds in the simultaneous presence of CRM1-RanGTP, CBC and capped RNA. As a consequence, both CRM1-RanGTP and the capped RNA are integrated into the complex with enhanced strength. In most structurally characterised CRM1-cargo complexes, CRM1 only specifically recognises the NES sequence. Within the snRNA export complex CRM1 is bound on multiple surfaces by the PHAX NES, PHAX^139-145^, PHAX^55-60^, ST2, and CBP20. The other well-documented case of extensive CRM1 recognition by a cargo is that of Snurportin1, which is involved in the nuclear import of snRNA^37^. However, the PHAX and Snurportin1 CRM1-binding surfaces, with the exception of their NESs, are not overlapping. The PHAX NES contains the conserved Q127 residue in the first position (Φ0), rather than a hydrophobic residue, indicative of a weak NES. In agreement, the Y154A mutant, designed not to be able to bind the RNA cap, still binds to CBC but not CRM1-RanGTP (Fig. 3h, lanes 7-9) indicating that the PHAX NES alone does not efficiently bind CRM1.

Finally, we investigated a potential role of ARS2 within the snRNA export complex. Since ARS2 also mediates degradative pathways, we wondered whether, how and at which point ARS2 might be excluded from the committed snRNA export complex. We have previously shown that the ARS2 effector domain interacts with the ARM motif of PHAX and also binds with its C-terminus to the groove at the CBP20-CBP80 interface^7,9^. *In vitro*, ARS2 prevents PHAX from directly binding to CBC in the CBCAP complex^7^. We now show that, in the presence of PHAX and CRM1-RanGTP, the CBC interacts more efficiently with PHAX, leading to a loss of the ARS2-CBP20-CBP80 groove interaction. In this context, ARS2 likely remains linked only via the PHAX ARM motif. Consequently, ARS2 might leave the complex and associate with other CBC-effector bound transcripts. In the case of ZC3H18, both the ARM-mediated interaction with ARS2 and Trp-helix binding to CBC are required for the activity of the NEXT complex^7^. The human/mouse PHAX ARM motif (9-EDGQL-13) is unique in comparison to the consensus sequence ([DE][DE]G[DE][ILV]), as it contains an unusual glutamine in the fourth position. *In vitro*, this exhibits lower affinity for ARS2 than ZC3H18, which has three tandem ARM motifs^7^. Whilst our *in vivo* analysis confirms that the PHAX ARM mutant variant does not interact with ARS2, such abolished interaction does not have an apparent impact on PHAX’s ability to recruit all snRNA export complex subunits (Figure 5k). This suggests that, in contrast to ZC3H18, ARS2 is not strictly required for the recruitment of PHAX to CBC-bound transcripts. In the absence of a robust ARS2 interaction, PHAX possibly evolved a stronger reliance on the CBC connection.

Efficient sorting between mRNAs and short transcripts, such as snRNAs, into their respective nuclear export pathways, as well as between useful and aberrant transcripts, requires RNA effectors. PHAX is believed to be only retained on short transcripts such as snRNAs, being competed out from longer transcripts >200–300 nts, such as most mRNAs, by hnRNP C ^20,25^. However it remains unclear exactly how this competition operates. Recently, the RNA helicase UAP56/DDX39B and ALYREF, which are subunits of the transcription-export (TREX) complex usually associated with mRNA export, were shown to stimulate the RNA binding of PHAX and participate in snRNA export^22^. How all these factors regulate nuclear export of different RNAs and the exact role of the PHAX-ARS2 interaction in this mechanism remains to be characterised.

## Methods

### Protein expression and purification

Human CBC was reconstituted as described previously^9^. Briefly, CBP80ΔNLS (lacking the N-terminal 19 residues) was expressed in High Five insect cells. CBP20 was expressed in *E. coli* BL21Star (DE3, Invitrogen) from the pETM30 vector (Gunter Stier, EMBL Heidelberg) as His-GST fusion. The cell pellets were lysed together in a buffer containing 20 mM Tris pH 8, 100 mM NaCl and 3% glycerol. Lysates were clarified by centrifugation and the complex was purified on a Ni^2+^-Chelating Sepharose (Cytiva). The His-GST tag was cleaved off by TEV protease. Following a subsequent passage through a Ni^2+^-Chelating Sepharose, CBC was loaded on a Heparin HiTrap column (Cytiva) and further purified by a gel filtration on Superdex 200 Increase (Cytiva).

Human ARS2^147-871^ and FL-PHAX were expressed as His-tag fusions in *E. coli* BL21Star (DE3, Invitrogen) from the pETM11 vector (Gunter Stier, EMBL Heidelberg). The proteins were purified on a Ni^2+^-chelating Sepharose (Cytiva) followed by HiTrap Heparin column (Cytiva) and gel filtration on Superdex 200 Increase (Cytiva) in a buffer containing 20 mM Tris pH 8, 100 mM NaCl, 3% glycerol^7^.

Human CRM1 was expressed as His-tag fusion protein from a pQE60 plasmid (Qiagen) in *E. coli* strain Tg1; and Human Ran containing the Q69L mutation was expressed as His-tag fusion from a pPROEX-HTB vector in *E. coli* BL21Star (DE3, Invitrogen). Harvested cells were resuspended in 20 mM Tris pH 8, 150 mM NaCl, 10 mM MgCl_2_, 3% glycerol and 0.5mM TCEP and lysed by sonication. Proteins were purified on Ni^2+^-chelating Sepharose (Cytiva) followed by Hitrap Heparin column (Cytiva). Proteins were further purified by gel filtration on Superdex 200 Increase (Cytiva) in a buffer containing 20 mM Tris pH 8, 100 mM NaCl, 2% glycerol.

PHAX^103-196^ and its mutant variants domains were expressed in *E. coli* BL21Star (DE3, Invitrogen) from the pETM41 vector (Gunter Stier, EMBL Heidelberg) as His-MBP fusion. The proteins were purified on a Ni^2+^-chelating Sepharose (Cytiva) followed by gel filtration on Superdex 200 Increase (Cytiva) in a buffer containing 20 mM Tris pH 8, 100 mM NaCl, 2% glycerol^7^.

### RNA preparation

U1SnRNA was transcribed in a 250 μl volume containing 12.5 μg of T7 RNA Polymerase (Promega), 10 mM DTT, 10 U RNasin Plus (Promega), NTP mixture (1 mM ATP, 1 mM CTP, 1 mM UTP and 1 mM GTP and 4 mM ARCA (Anti-Reverse Cap Analog) (NEB)), 3 μg DNA template, 1mM spermidine, 15 mM MgCl_2_ and 50 mM Tris pH8, for 3h at 37°C. Sample was treated with 40U of DNAse I for 10min at 37°C. RNA was purified using Zymo R1017 columns following manufacturer instructions. Transcription product was analysed on SDS-PAGE. RNA was refolded at 95°C for 5 min followed by cool-down at room temperature for 1h in presence of 1mM MgCl_2._

### Protein Phosphorylation

100 μg of PHAX purified protein was incubated for 1h at room temperature in presence of 1000 U of CK2, 100 μM ATP and 2.5 μl reaction Buffer (NEB).

### RanGTP/GDP exchange

Typically, 30 nmol of Ran were incubated with 10 mM EDTA and 7% phosphatase alkaline at room temperature for 2h. Sample was spined at 2000g for 10 min and purified on Superdex 200 Increase (Cytiva) in a buffer containing 20 mM Tris pH 8, 100 mM NaCl, 2% glycerol. Protein sample was mixed with 1mM GTP and purified on Superdex 200 Increase (Cytiva) in a buffer containing 20mM Tris pH 8, 100 mM NaCl, 2% glycerol and 50 μM GTP, flash frozen in liquid nitrogen and stored at -80°C.

### Assembly of the snRNA export complex

4 nmol of CBC and 8nmol of phosphorylated PHAX, CRM1 and RanGTP were mixed in presence of 100μM U1 SnRNA or short capped RNA (m^7^GpppAAUCUAUAAUAGCA), 1 mM Mg^2+^, 1mM KCl and 100 μM GTP. Complex was loaded on Superdex 200 Increase 3.2/300 with 100 mM NaCl, 20 mM NaCl, 1 mM Mg^2+^, 0.5 mM TCEP, 2% glycerol and 10 μM GTP. Purified sample was used directly for cryo-EM grid preparation.

### Complex assembly analysis by gel filtration

180pmol of CBC and 360pmol of the different PHAX constructs, CRM1 and RanGTP were mixed in presence of 100μM capped-RNA (m^7^GpppAAUCUAUAAUAGCA), cap-analogue m^7^GpppG or m^7^-GTP and 1mM Mg^2+^, 1mM KCl, 100μM GTP. Samples were incubated for 30min on ice before analysis on Superdex 200 Increase 3.2/300. 50μl fractions were collected and analysed on SDS-PAGE.

### Isothermal Calorimetry (ITC)

ITC experiments were performed at 25°C using a PEAQ-ITC micro-calorimeter (MicroCal). Experiments included one 0.5 µl injection and 15-18 injections of 1.5µL at 0.2-0.25 mM of m^7^GTP, m^7^GpppG or PHAX^103-196^ and its R147E, Y152E, Y154E mutant into the sample cell that contained 20µM CBC in presence or absence of 0.1 mM m^7^GTP or m^7^GpppG, in 20 mM Tris pH 8.0, 100 mM NaCl, 2%glycerol. The initial data point was deleted from the data sets. Binding isotherms were fitted with a one-site binding model by nonlinear regression using the MicroCal PEAQ-ITC Analysis software. Measurements were performed in duplicate.

### Mass photometry analysis

Mass photometry measurements were performed on a OneMP mass photometer (Refeyn). Coverslips (No. 1.5H, 24 × 50 mm, VWR) were washed with water and isopropanol before being used as a support for silicone gaskets (CultureWellTM 423 Reusable Gaskets, Grace Bio-labs). Contrast/mass calibration was realised using native marker (Native Marker unstained protein 426 standard, LC0725, Life Technologies) with a medium field of view and monitored during 60 s using the AcquireMP software (Refeyn). For each condition, 18 µl of buffer 100 mM NaCl, 20 mM NaCl, 1 mM Mg^2+^, 0.5 mM TCEP, 2% glycerol and 10 μM GTP were used to find the focus. 2 µl of diluted purified complex or diluted equimolar mix of the different proteins and RNA (m^7^GpppAAUCUAUAAUAGCA) (200 nM) were added to reach a final concentration of 20 nM. Movies of 60 s were recorded, processed and mass estimation was determined automatically using the DiscoverMP software (Refeyn).

### Cell culture and transfections

All mouse embryonic stem (mES) cell lines used or generated in this study were descendants of wildtype ES-E14TG2a cells (male genotype, XY). mES cells were cultured on 0.2% gelatine coated plates in 2i/LIF containing media (1:1 mix of DMEM/F12 (Thermo Fisher,) and Neurobasal (Thermo Fisher) supplemented with 1% (v/v) penicillin-streptomycin (Sigma,), 2 µM Glutamax (Thermo Fisher), 0.1 mM Non-Essential Amino Acids (Thermo Fisher), 1 mM Sodium Pyruvate (Thermo Fisher), 0.5x N-2 Supplement (Thermo Fisher), 0.5x B-27 Supplement (Thermo Fisher), 50 µM 2-Mercaptoethanol (Thermo Fisher) 3 µM GSK3- inhibitor (CHIR99021, Sigma), 1 µM MEK-inhibitor (PD0325901, Sigma) and Leukemia Inhibitory Factor (LIF, produced in house)) at 37°C, 5% CO₂. Cells were passaged every 48- 72 hours by dissociating cells with 0.05% Trypsin (Sigma) before neutralizing with an equal volume of 1x Trypsin Inhibitor (Thermo Fisher). Cells were pelleted by centrifugation to remove Trypsin before resuspending in 2i/LIF media and plating at ∼ 8x10^4^ cells/ml.

The generation of CRISPR/Cas9-mediated genomic knock-ins of N-terminal 2xHA- FKBP-V(HA-dTAG)-PHAX mES cells was carried out as described in^49^. Depletion of HA-dTAG-PHAX was performed by the addition of dTAG^V^-1 (Tocris) to the cell culture medium to a final concentration of 100 nM.

### cDNA cloning and exogenous expression of PHAX

Full length mouse PHAX (mmPHAX) cDNA was cloned from cDNA libraries synthesis from 2 µg total mouse RNA using Superscript III reverse transcription reagents (Thermo Fisher) into a pCR8/GW/TOPO Gateway entry plasmid (Thermo Fisher) by TA TOPO cloning. PHAX mutations were introduced into pCR8[mmPHAX] plasmids by PCR. These were then used as templates to introduce into a piggyBAC (pB) vector containing an N-terminal MYC and miniTurbo (MYC-mTurbo) tag using NEBbuilder HiFi DNA assembly (NEB). Wildtype mES cells were transfected with pB[MYC-] or pB[MYC-mTurbo-PHAX] vectors along with a piggyBAC transposase expressing vector (pBase) in a 1:1 ratio using Viafect (Promega). Cell pools were selected for successful integration of pB plasmids with Blastocidin (BSD) for 7-10 days until negative control cells no longer survived. Expression of constructs were validated by western blotting analysis using MYC antibodies.

### *In vivo* proximity labelling and streptavidin enrichment

*In vivo* proximity labelling (PL) with MYC-mTurbo-PHAX variants was stimulated by supplementing the media with 50 µM biotin for 4 hours. Cells expressing MYC without the mTurbo fusion were used in parallel as negative controls. For the enrichment of biotinylated proteins, cells were lysed in HT150 extraction buffer (20 mM HEPES pH 7.4, 150 mM NaCl, 0.5% v/v Triton X-100) freshly supplemented with protease inhibitors. Lysates were sheared by sonication (3 x 5 s, amplitude 2) and cleared by centrifugation at 18,000g for 20 minutes. Biotinylated proteins were enriched by incubating with Dynabeads MyOne Streptavidin C1 beads (ThermoFisher) overnight at 4°C. Beads were washed 3 times with HT150 extraction buffer, transferring beads to a fresh tube on the final wash. Proteins were eluted by boiling in 1X NuPAGE loading buffer (Invitrogen) for 5 minutes. Supernatants were mixed with 10X Reducing Agent (Invitrogen) and denatured for a further 5 minutes at 95°C before proceeding with western blotting analysis. To assess biotinylation, membranes were first blocked in 10% w/v BSA before probing with HRP-Conjugated Streptavidin (ThermoFisher). Bands were visualized by SuperSignal West Femto chemiluminescent ECL (Thermo) and captured using an Amersham Imager 600 or ImageQuant 800 imagine systems (GE Healthcare).

### Cryo-EM sample preparation

EM grids were glow-discharged on each side for 90s at 30 sccm, with 100% power with a mixture of 90% argon and 10% oxygen using the Fischione 1070 Nanoclean plasma cleaner. 3.5 μl of the sample was applied to 300 mesh Quantifoil R 1.2/1.3 glow-discharged grids, at 4 °C, 100 % humidity and blotting for 2s at blot force -2 in a Vitrobot Mark IV.

### Cryo-EM data collection and processing

Grids were initially screened using a Glacios Cryo-TEM equipped with a Falcon 4i Direct Electron Detector and SelectrisX energy filter (Thermo Fisher Scientific). Final data collection was performed on a Titan Krios TEM (Thermo Fisher Scientific) operated at 300 kV equipped with a Gatan K3 and a GIF Quantum energy filter (Gatan). 8000 movies were collected at 105k magnification giving a pixel size of 0.84 Å, a defocus range from -1.0 to -2.2 µm in 0.2 µm step and a total dose of ∼40 e-/ Å^2^ per movie and 40 frames per movie.

Data processing was performed using an cryoSPARC v4.3^50^. After Movie drift correction and CTF parameters determination, realigned micrographs were manually inspected resulting in 7302 selected micrographs. Particles were firstly picked on 665 representative micrographs using Blob Picker with a diameter ranging from 110 to 190 Å and then extracted with a box size of 400-400 Å. The best particles were used to generate a Topaz model or to create Templates for particles picking. After extraction, duplicates were removed and 2D classification was applied to eliminate particles displaying poor structural features and to select 2D class averages with lower background noise. ‘Ab-initio’ reconstruction was performed, followed by ‘heterogeneous refinement’ and ‘3D classification’ to further select particles. ‘Non-uniform refinement’ was performed on the final sets of particles, followed by ‘reference based motion correction’ and a final ‘Non-uniform refinement’. The final map was at an average resolution of 2.62 Å (FSC 0.143 threshold) (Supplementary Table 1). Based on this consensus map, particle subtraction around CBC and CRM1/RanGTP was performed. The subtracted particles were finally subject to local refinement to improve subtracted particle angles and shifts estimation. All final maps were used to calculate directional FSC and local resolution in CryoSPARC. The detailed processing workflow, comprising data statistics, is shown in Supplementary Fig. 2.

### Model building and refinement

Model building, using available structures of CBC-cap and CRM1-RanGTP as a guide, was performed in COOT^51^ initially using the high-resolution maps focussed on the CBC or CRM1-RanGTP moieties of the snRNA export complex. The AF2 prediction of the complete complex was also useful. Since PHAX is at the slightly flexible interface between the two lobes of the complex, final refinement of the complete complex was against the composite focussed map. Models were refined using Phenix real-space refinement^52^ with Ramachandran restraints (Supplementary Table 1). Structure figures were prepared using ChimeraX^53^. Electrostatic potential was calculated using ChimeraX^53^.

## Data Availability

The atomic coordinates and cryo-EM maps of the human snRNA export complex determined in this study have been deposited under the PDB accession code 9HFL and EMDB code 52115, respectively.

## Acknowledgements

SC and JK were funded by the ANR MTREC grant (ANR-21-CE11-0021). THJ was supported by the Danish National Research Council and the Novo Nordisk Foundation [ExoAdapt Grant 31199]. IBS acknowledges integration into the Interdisciplinary Research Institute of Grenoble (IRIG, CEA). This work used the platforms of the Grenoble Instruct-ERIC center (ISBG ; UAR 3518 CNRS-CEA-UGA-EMBL) within the Grenoble Partnership for Structural Biology (PSB), supported by FRISBI (ANR-10-INBS-0005-02) and GRAL, financed within the University Grenoble Alpes graduate school (Ecoles Universitaires de Recherche) CBH-EUR-GS (ANR-17-EURE-0003). We thank Caroline Mas for assistance with ITC and mass photometry. We thank Wojtek Galej, Iskander Khusainov and Romain Linares for access to the Glacios at EMBL Grenoble. We acknowledge the European Synchrotron Radiation Facility for provision of beam time on CM01 and we would like to thank Gregory Effantin for assistance^54^. We thank Anne-Emmanuelle Foucher for help with protein purification. We thank Nadia Laurs Schmidt for excellent technical assistance.

## Author contributions

E.D. performed all *in vitro* biochemical and biophysical characterization and cryo-EM sample preparation. E.D. and S.C. performed cryo-EM structural characterization. M.F.B., W.G. and T.H.J. performed all the cellular assays. E.D., J.K and S.C conceived the project. J.K and S.C wrote the manuscript with input from all authors.

---

**Supplementary Figure 1.**
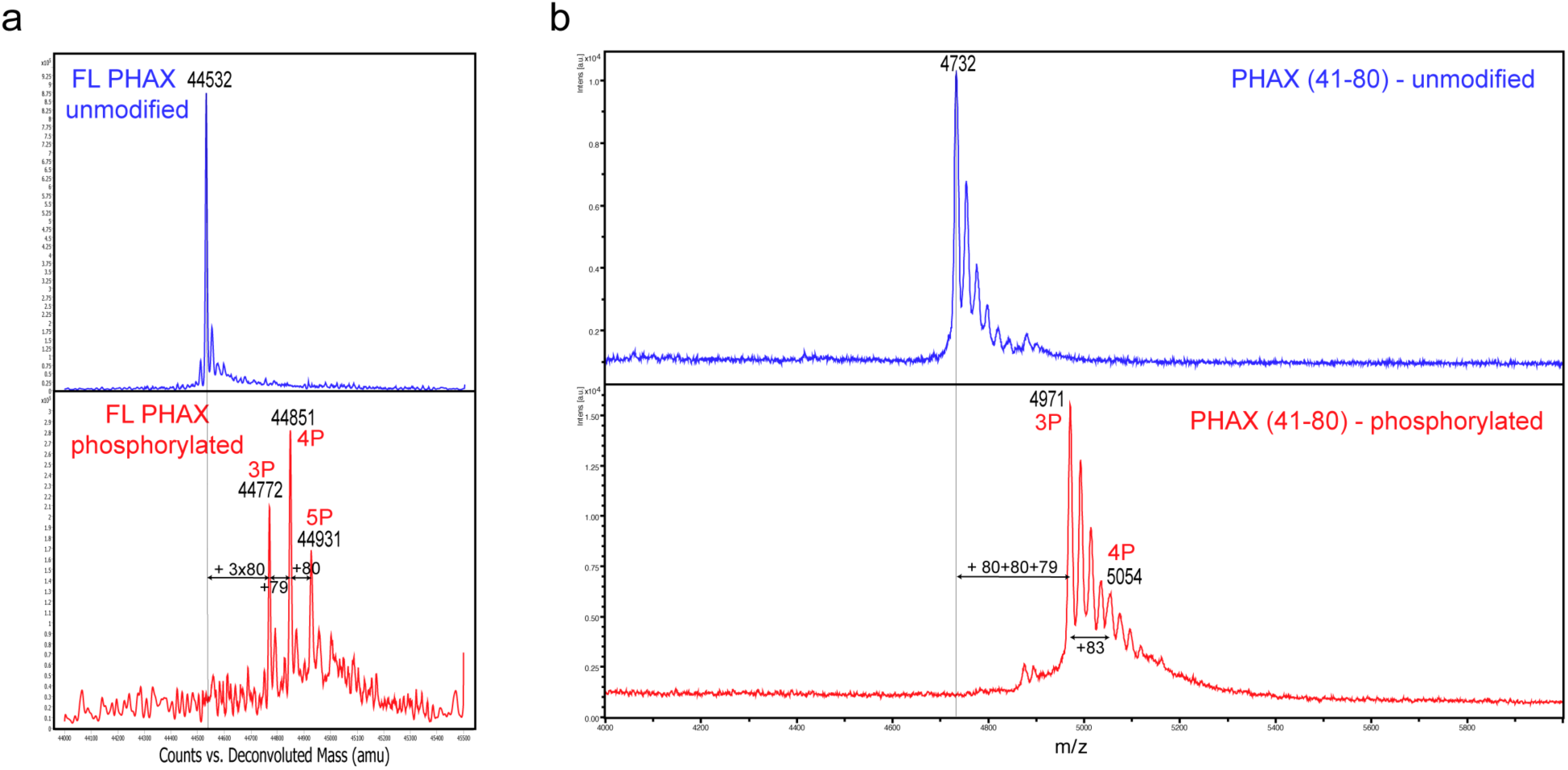
PHAX phosphorylation analysis. Masses of CK2-treated FL PHAX and PHAX^41-80^ were assessed using mass spectrometry (MS). **a.** FL PHAX was analysed by LC-ESI-MS (liquid chromatography-electrospray). Data are presented after deconvolution of *m/z* (mass-to-charge ratio) to masses (expressed in Da). Spectra of FL PHAX before and after phosphorylation were compared. The full- length protein has between 3 and 5 phosphorylated sites (3P, 4P and 5P). b) PHAX (41-80) was analysed by MALDI-MS (Matrix Assisted Laser Desorption Ionisation-mass spectrometry). MALDI-MS determines *m/z* values of singly charged PHAX^41-80^ ions. The *m/z* difference between the untreated PHAX (41-80) and the phosphorylated one indicates that PHAX^41-80^ carries 3 phosphorylated sites (3P). An additional site (4P) may be present. Peaks that are not labelled are due to adducts with non-volatile molecules (such as Na).

**Supplementary Figure 2.**
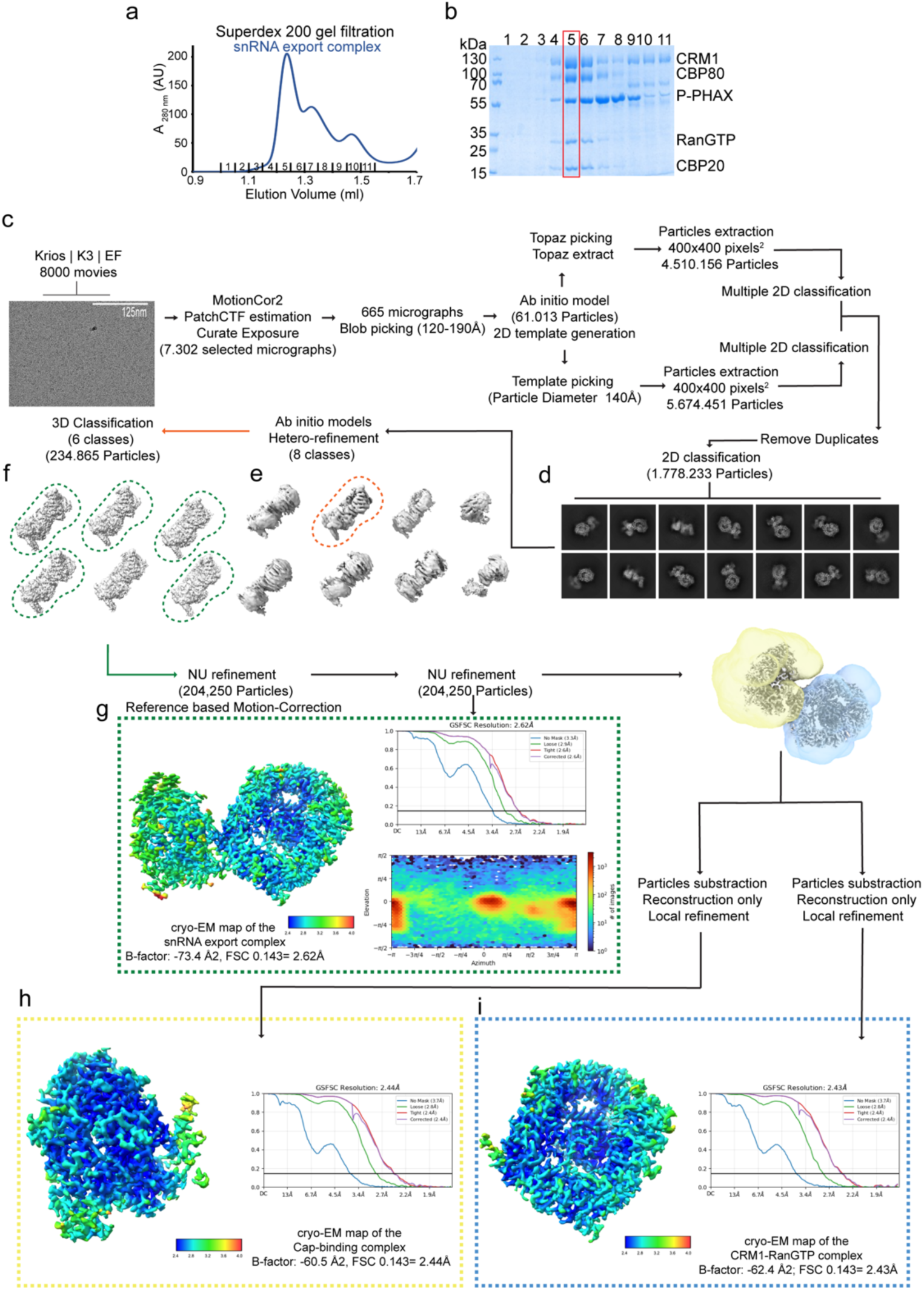
Cryo-EM image processing strategy used to obtain the structure of the snRNA export complex. **a,b.** Superdex 200 gel filtration elution profile and SDS-PAGE analysis of fraction 1-12 of the snRNA complex reconstituted with capped 14nt RNA. The complex eluted in fraction 5 used for grid preparation is highlighter with a red rectangle in **b**. **c.** Representative cropped micrograph (Scale bar = 150nm). **d.** 2D class averages. **e,f.** 3D class averages. **g-i.** Views of each local resolution sharpened EM maps are shown. Fourier shell correlation curves (FSC) are displayed.

**Supplementary Figure 3.**
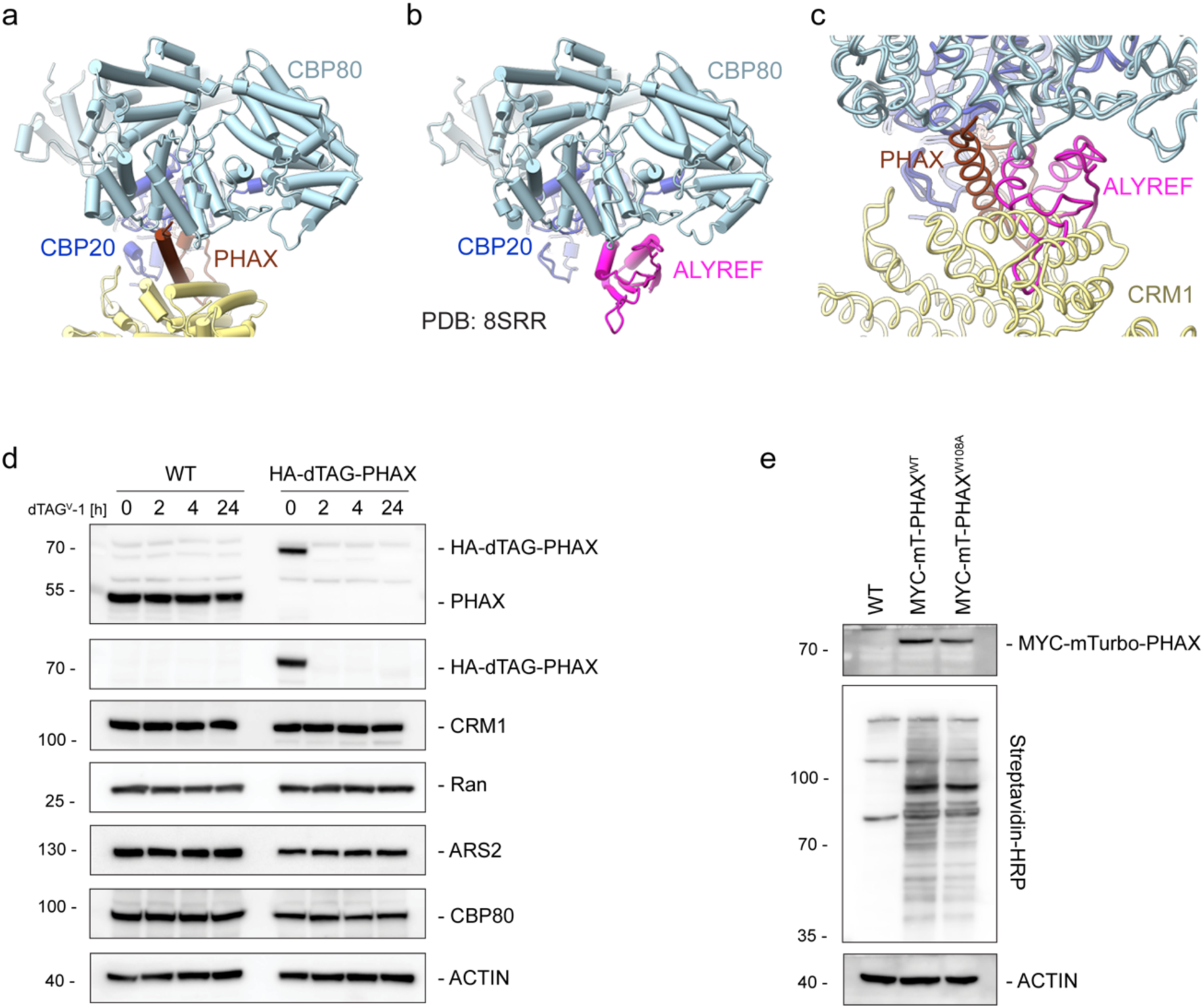
CBC Binding of ALYREF is incompatible with binding of PHAX and CRM1. **a-c.** Comparison of CBC interactions with PHAX and CRM1 (**a**) with the CBC contacts with ALYREF (**b**) (PDB:8SRR, shown in magenta). The two structures were superimposed using CBC (**c**). The binding of ALYREF to the CBC is incompatible with that of PHAX and CRM1. **(d)** Western blot analysis of lysates from WT or HA-dTAG-PHAX mES cells harvested at indicated time points (hours) following dTAG^V^-1 treatment. Membranes were probed with antibodies against PHAX, HA, CRM1, Ran, ARS2 and CBP80. ACTIN was used as a loading control. **(e)** Western blot analysis of lysates from WT control or HA-dTAG- PHAX mES cells expressing MYC-miniTurbo(mT)-PHAX proteins. Cells were treated with dTAG^V^-1 to deplete endogenous dTAG-PHAX and biotin was added for 6 hours to induce proximity labelling. Membranes were probed with MYC and ACTIN to assess protein expression and Streptavidin-HRP to assess biotinylation efficiency of mTurbo-tagged proteins.

**Supplementary Figure 4.**
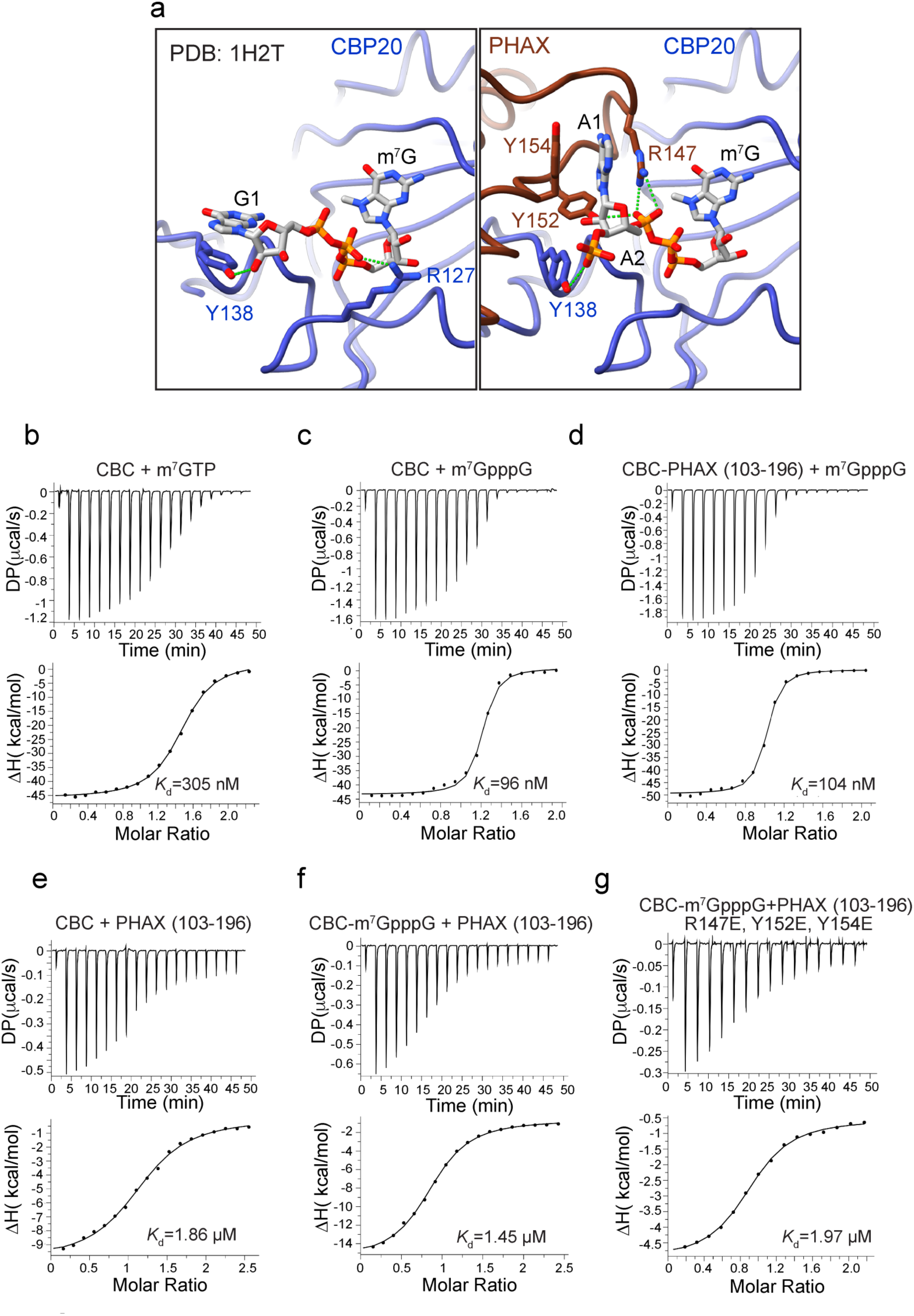
Characterization of RNA recognition by PHAX. **a.** Comparison of the molecular details of RNA cap binding by CBC and CBC-PHAX. **b.** ITC measurement of the interaction affinity between CBC and of m^7^GTP. **c,d.** ITC measurements of the interaction affinity between CBC and cap analogue (m^7^GpppG) in absence or presence of PHAX^103-196^. **e,f.** ITC measurements of the interaction affinity between CBC and PHAX^103-196^ in absence or presence of m^7^GpppG. **g.** ITC measurement of the interaction affinity between CBC-m^7^GpppG and the R147E, Y152E, Y154E mutant of PHAX^103-196^_._

**Supplementary Figure 5.**
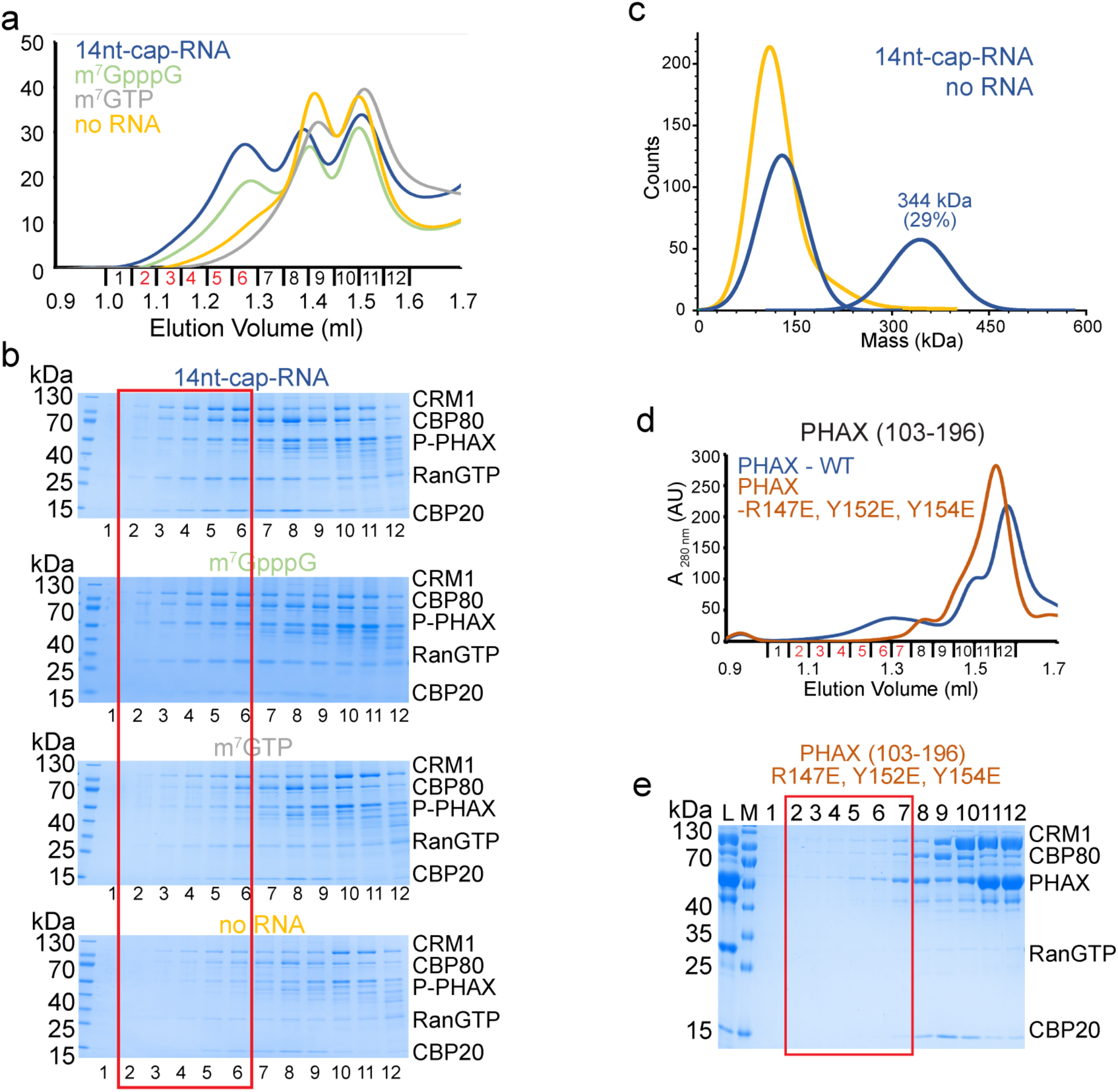
RNA is required for snRNA complex formation. **a.** Overlay of Superdex 200 gel filtration elution profiles of the snRNA complex reconstitutions with different lengths of RNA. **b.** SDS-PAGE analysis of fractions 1-12 of the Superdex 200 gel filtration elution profiles shown in **a**. **c.** Mass photometry analysis of the snRNA complex formed with phosphorylated FL PHAX in the presence or absence of 14nt capped RNA. The measured molecular mass the full complex and the percentage of corresponding observations is indicated. **d.** Overlay of Superdex 200 gel filtration elution profiles of the snRNA complex reconstitutions with using either WT or the R147E, Y152E, Y154E mutant of MBP-PHAX^103-196^ and a short capped 14nt RNA. The WT complex elutes in fractions highlighted in red. **d.** SDS-PAGE analysis of fractions 1-12 of the gel filtration elution profile shown in **c.** MBP- PHAX^103-196^ R147E, Y152E, Y154E is unable to form the complex with CBC and CRM1-RanGTP. The fractions corresponding to the WT complex elution peak are highlighted with red rectangle.

**Supplementary Figure 6.**
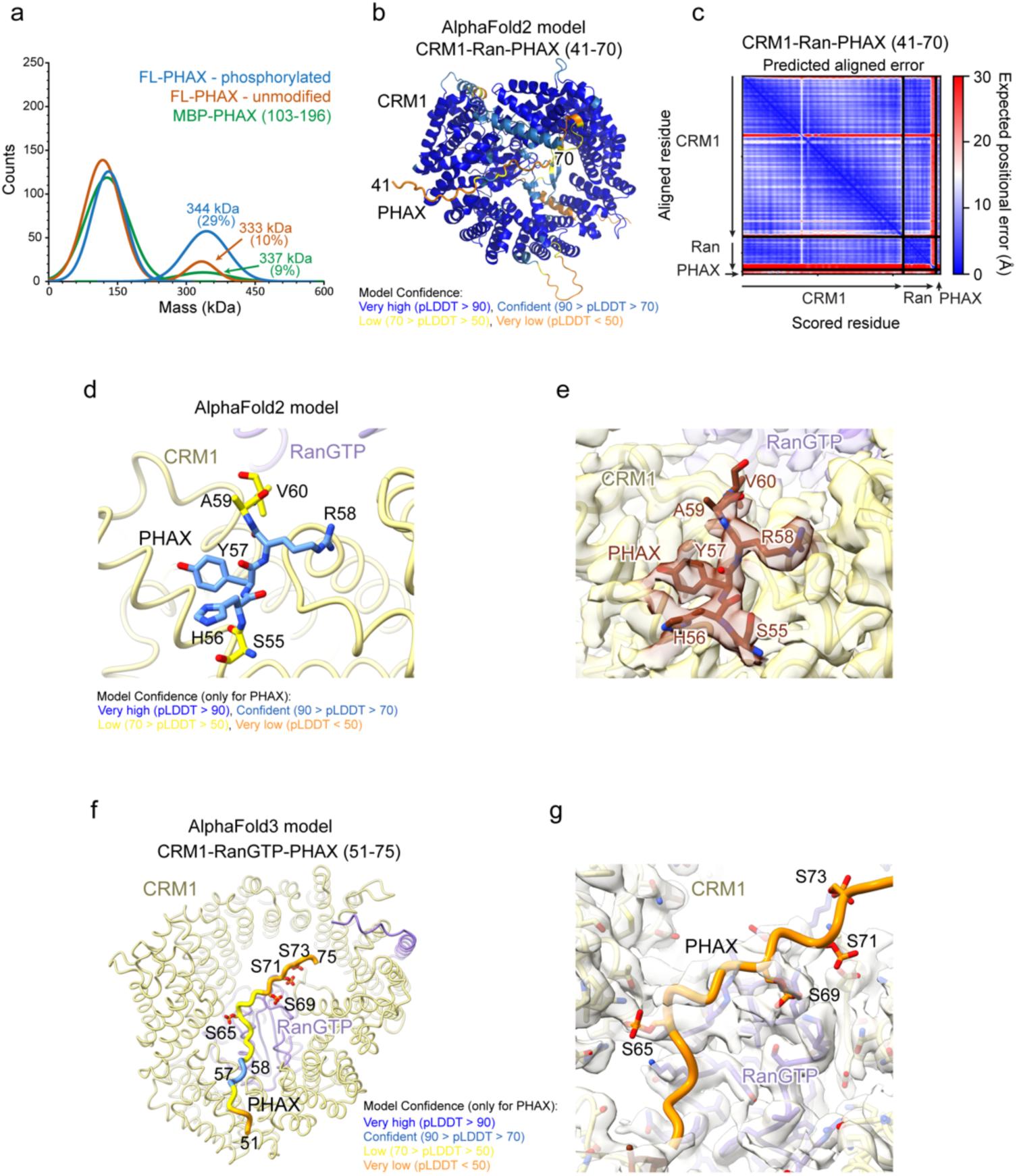
CRM1-Ran-PHAX structure prediction. **c.** Mass photometry analysis of the snRNA complex formed with phosphorylated FL PHAX, unmodified FL PHAX and MBP-PHAX^103-196^ in the presence of 12nt capped RNA. The measured molecular mass of the full complex and the percentage of corresponding observations is indicated. **b.** AlphaFold2 predicted structure of CRM1-Ran-PHAX^41-70^ coloured according to pLDDT. **c.** Predicted aligned error (PAE) plot for the CRM1-Ran-PHAX^41-70^ model. **d.** AlphaFold2 model of CRM1-Ran-PHAX^41-70^. Only residues 55- 60 of PHAX are shown as sticks. Only PHAX is coloured according to pLDDT. CRM1 is shown in yellow. **e.** Cryo-EM map and details of the interactions between CRM1 and the PHAX^56-60^. **f.** AlphaFold3 model of CRM1-Ran-PHAX^1-103^ with phosphorylation on serines 65,69,71 and 73. Only residues 55-75 of PHAX are shown. Only PHAX is coloured according to pLDDT. CRM1 is shown in yellow and Ran GTP in violet. **g.** Extra electron density, appearing at low contour level, that partially overlaps with positions of phosphorylated serines predicted by AlphaFold3.

**Supplementary Table 1.** Cryo-EM data collection, refinement and validation statistics.

| snRNA export complex<br>(PDB-9HFL, EMD-52115) |  |  |  |
| --- | --- | --- | --- |
| Data collection and processing |  |  |  |
| Date of data collection | 27-11-2023 |  |  |
| Microscope (ESRF) | Krios K2, EF |  |  |
| Magnification | 105,000 x |  |  |
| Voltage (kV) | 300 |  |  |
| Electron exposure (e-/Å<sup>2</sup>) | 40.5 |  |  |
| Defocus range (µm) | -1.0 and -2.2 |  |  |
| Pixel size (Å/pix) | 0.84 |  |  |
| SuperRes pixel size (Å/pix) | n/a |  |  |
| Symmetry imposed | - |  |  |
| Initial particle images (no.) | 1778233 |  |  |
| Final particle images (no.) | 204250 |  |  |
|  | Complete map | Focus map CBC | Focus map CRM1 |
| Global map resolution (Å) | 2.62 | 2.44 | 2.43 |
| 3DFSC resolution (Å) | 2.71 | 2.60 | 2.56 |
| FSC threshold 0.143 |  |  |  |
| Map resolution range (Å) (25<sup>th</sup> to 75<sup>th</sup> percentile) | 2.25-3.85 | 2.05-3.50 | 2.10-4.01 |
| Map sharpening *B* factor (Å<sup>2</sup>) | -73.4 | -60.5 | -62.4 |
| Refinement and Validation |  |  |  |
| Model composition |  |  |  |
| Non-hydrogen atom | 17802 |  |  |
| Protein residues | 2159 |  |  |
| Waters | 32 |  |  |
| Ligands | 3 (m7GpppA, GTP, Mg) |  |  |
| *B* factor (Å<sup>2</sup>) |  |  |  |
| Protein residues | 54.05 |  |  |
| Waters | 39.70 |  |  |
| Ligands | 39.30 |  |  |
| R.m.s. deviations |  |  |  |
| Bond lengths (Å) | 0.002 |  |  |
| Bond angles (°) | 0.428 |  |  |
| Validation |  |  |  |
| MolProbity score | 1.38 |  |  |
| Clashscore | 3.30 |  |  |
| Poor rotamers (%) | 2.19 |  |  |
| Ramachandran plot |  |  |  |
| Favored (%) | 98.09 |  |  |
| Allowed (%) | 1.91 |  |  |
| Disallowed (%) | 0.00 |  |  |
| Model vs Data |  |  |  |
| CC (mask) | 0.82 |  |  |
| CC (box) | 0.65 |  |  |
| CC (peaks) | 0.63 |  |  |
| CC (volume) | 0.81 |  |  |
| Mean CC for ligands | 0.67 |  |  |

## References

1. Carninci, P., Kasukawa, T., Katayama, S., Gough, J., Frith, M.C., Maeda, N., Oyama, R., Ravasi, T., Lenhard, B., Wells, C., et al. (2005). The Transcriptional Landscape of the Mammalian Genome. Science 309, 1559–1563. 10.1126/science.1112014.

2. Djebali, S., Davis, C.A., Merkel, A., Dobin, A., Lassmann, T., Mortazavi, A., Tanzer, A., Lagarde, J., Lin, W., Schlesinger, F., et al. (2012). Landscape of transcription in human cells. Nature 489, 101–108. 10.1038/nature11233.

3. Jensen, T.H., Jacquier, A., and Libri, D. (2013). Dealing with Pervasive Transcription. Molecular Cell 52, 473–484. 10.1016/j.molcel.2013.10.032.

4. Izaurralde, E., Lewis, J., McGuigan, C., Jankowska, M., Darzynkiewicz, E., and Mattaj, I.W. (1994). A nuclear cap binding protein complex involved in pre-mRNA splicing. Cell 78, 657–668. 10.1016/0092-8674(94)90530-4.

5. Rambout, X., and Maquat, L.E. (2020). The nuclear cap-binding complex as choreographer of gene transcription and pre-mRNA processing. Genes Dev. 34, 1113–1127. 10.1101/gad.339986.120.

6. Cho, H., Rambout, X., Gleghorn, M.L., Nguyen, P.Q.T., Phipps, C.R., Miyoshi, K., Myers, J.R., Kataoka, N., Fasan, R., and Maquat, L.E. (2018). Transcriptional coactivator PGC-1α contains a novel CBP80-binding motif that orchestrates efficient target gene expression. Genes Dev. 32, 555–567. 10.1101/gad.309773.117.

7. Dubiez, E., Pellegrini, E., Finderup Brask, M., Garland, W., Foucher, A.-E., Huard, K., Heick Jensen, T., Cusack, S., and Kadlec, J. (2024). Structural basis for competitive binding of productive and degradative co-transcriptional effectors to the nuclear cap-binding complex. Cell Reports 43, 113639. 10.1016/j.celrep.2023.113639.

8. Mazza, C., Segref, A., Mattaj, I.W., and Cusack, S. (2002). Large-scale induced fit recognition of an m7GpppG cap analogue by the human nuclear cap-binding complex. The EMBO Journal 21, 5548–5557. 10.1093/emboj/cdf538.

9. Schulze, W.M., and Cusack, S. (2017). Structural basis for mutually exclusive co-transcriptional nuclear cap-binding complexes with either NELF-E or ARS2. Nat Commun 8, 1302. 10.1038/s41467-017-01402-w.

10. Calero, G., Wilson, K.F., Ly, T., Rios-Steiner, J.L., Clardy, J.C., and Cerione, R.A. (2002). Structural basis of m7GpppG binding to the nuclear cap-binding protein complex. Nat Struct Biol 9, 912–917. 10.1038/nsb874.

11. Lykke-Andersen, S., Rouvière, J.O., and Jensen, T.H. (2021). ARS2/SRRT: at the nexus of RNA polymerase II transcription, transcript maturation and quality control. Biochemical Society Transactions 49, 1325–1336. 10.1042/BST20201008.

12. Garland, W., and Jensen, T.H. (2024). Nuclear sorting of short RNA polymerase II transcripts. Molecular Cell 84, 3644–3655. 10.1016/j.molcel.2024.08.024.

13. Foucher, A.-E., Touat-Todeschini, L., Juarez-Martinez, A.B., Rakitch, A., Laroussi, H., Karczewski, C., Acajjaoui, S., Soler-López, M., Cusack, S., Mackereth, C.D., et al. (2022). Structural analysis of Red1 as a conserved scaffold of the RNA-targeting MTREC/PAXT complex. Nat Commun 13, 4969. 10.1038/s41467-022-32542-3.

14. Polák, P., Garland, W., Rathore, O., Schmid, M., Salerno-Kochan, A., Jakobsen, L., Gockert, M., Gerlach, P., Silla, T., Andersen, J.S., et al. (2023). Dual agonistic and antagonistic roles of ZC3H18 provide for co-activation of distinct nuclear RNA decay pathways. Cell Reports 42, 113325. 10.1016/j.celrep.2023.113325.

15. Dobrev, N., Ahmed, Y.L., Sivadas, A., Soni, K., Fischer, T., and Sinning, I. (2021). The zinc-finger protein Red1 orchestrates MTREC submodules and binds the Mtl1 helicase arch domain. Nat Commun 12, 3456. 10.1038/s41467-021-23565-3.

16. Staněk, D. (2017). Cajal bodies and snRNPs - friends with benefits. RNA Biology 14, 671–679. 10.1080/15476286.2016.1231359.

17. Suzuki, T., Izumi, H., and Ohno, M. (2010). Cajal body surveillance of U snRNA export complex assembly. Journal of Cell Biology 190, 603–612. 10.1083/jcb.201004109.

18. Heath, C.G., Viphakone, N., and Wilson, S.A. (2016). The role of TREX in gene expression and disease. Biochemical Journal 473, 2911–2935. 10.1042/BCJ20160010.

19. Hohmann, U., Graf, M., Schellhaas, U., Pacheco-Fiallos, B., Fin, L., Riabov-Bassat, D., Pühringer, T., Szalay, M.-F., Tirián, L., Handler, D., et al. (2024). A molecular switch orchestrates the nuclear export of human messenger RNA. bioRxiv 2024.03.24.586400; doi: 10.1101/2024.03.24.586400.

20. Ohno, M., Segref, A., Bachi, A., Wilm, M., and Mattaj, I.W. (2000). PHAX, a Mediator of U snRNA Nuclear Export Whose Activity Is Regulated by Phosphorylation. Cell 101, 187–198. 10.1016/S0092-8674(00)80829-6.

21. Izumi, H., McCloskey, A., Shinmyozu, K., and Ohno, M. (2014). p54nrb/NonO and PSF promote U snRNA nuclear export by accelerating its export complex assembly. Nucleic Acids Research 42, 3998–4007. 10.1093/nar/gkt1365.

22. Taniguchi, I., Hirose, T., and Ohno, M. (2024). The RNA helicase DDX39 contributes to the nuclear export of spliceosomal U snRNA by loading of PHAX onto RNA. Nucleic Acids Research, gkae622. 10.1093/nar/gkae622.

23. Dickmanns, A., Monecke, T., and Ficner, R. (2015). Structural Basis of Targeting the Exportin CRM1 in Cancer. Cells 4, 538–568. 10.3390/cells4030538.

24. Kitao, S., Segref, A., Kast, J., Wilm, M., Mattaj, I.W., and Ohno, M. (2008). A Compartmentalized Phosphorylation/Dephosphorylation System That Regulates U snRNA Export from the Nucleus. Molecular and Cellular Biology 28, 487–497. 10.1128/MCB.01189-07.

25. McCloskey, A., Taniguchi, I., Shinmyozu, K., and Ohno, M. (2012). hnRNP C Tetramer Measures RNA Length to Classify RNA Polymerase II Transcripts for Export. Science 335, 1643–1646. 10.1126/science.1218469.

26. Ohno, M. (2012). Size matters in RNA export. RNA Biol 9, 1413–1417. 10.4161/rna.22569.

27. Dantsuji, S., Ohno, M., and Taniguchi, I. (2023). The hnRNP C tetramer binds to CBC on mRNA and impedes PHAX recruitment for the classification of RNA polymerase II transcripts. Nucleic Acids Research 51, 1393–1408. 10.1093/nar/gkac1250.

28. Xie, M., Li, M., Vilborg, A., Lee, N., Shu, M.-D., Yartseva, V., Šestan, N., and Steitz, J.A. (2013). Mammalian 5′-Capped MicroRNA Precursors that Generate a Single MicroRNA. Cell 155, 1568– 1580. 10.1016/j.cell.2013.11.027.

29. Machitani, M., Taniguchi, I., McCloskey, A., Suzuki, T., and Ohno, M. (2020). The RNA transport factor PHAX is required for proper histone H2AX expression and DNA damage response. RNA 26, 1716–1725. 10.1261/rna.074625.120.

30. Mourão, A., Varrot, A., Mackereth, C.D., Cusack, S., and Sattler, M. (2010). Structure and RNA recognition by the snRNA and snoRNA transport factor PHAX. RNA 16, 1205–1216. 10.1261/rna.2009910.

31. Segref, A., Mattaj, I.W., and Ohno, M. (2001). The evolutionarily conserved region of the U snRNA export mediator PHAX is a novel RNA-binding domain that is essential for U snRNA export. RNA 7, 351–360. 10.1017/S1355838201002278.

32. Hallais, M., Pontvianne, F., Andersen, P.R., Clerici, M., Lener, D., Benbahouche, N.E.H., Gostan, T., Vandermoere, F., Robert, M.-C., Cusack, S., et al. (2013). CBC–ARS2 stimulates 3′-end maturation of multiple RNA families and favors cap-proximal processing. Nat Struct Mol Biol 20, 1358–1366. 10.1038/nsmb.2720.

33. Schulze, W.M., Stein, F., Rettel, M., Nanao, M., and Cusack, S. (2018). Structural analysis of human ARS2 as a platform for co-transcriptional RNA sorting. Nat Commun 9, 1701. 10.1038/s41467-018-04142-7.

34. Cook, A.G., Fukuhara, N., Jinek, M., and Conti, E. (2009). Structures of the tRNA export factor in the nuclear and cytosolic states. Nature 461, 60–65. 10.1038/nature08394.

35. Okada, C., Yamashita, E., Lee, S.J., Shibata, S., Katahira, J., Nakagawa, A., Yoneda, Y., and Tsukihara, T. (2009). A high-resolution structure of the pre-microRNA nuclear export machinery. Science 326, 1275–1279. 10.1126/science.1178705.

36. Smith, A.M., Li, Y., Velarde, A., Cheng, Y., and Frankel, A.D. (2024). The HIV-1 Nuclear Export Complex Reveals the Role of RNA in Crm1 Cargo Recognition. https://www.biorxiv.org/content/10.1101/2024.09.22.614349v1.

37. Monecke, T., Güttler, T., Neumann, P., Dickmanns, A., Görlich, D., and Ficner, R. (2009). Crystal Structure of the Nuclear Export Receptor CRM1 in Complex with Snurportin1 and RanGTP. Science 324, 1087–1091. 10.1126/science.1173388.

38. Güttler, T., Madl, T., Neumann, P., Deichsel, D., Corsini, L., Monecke, T., Ficner, R., Sattler, M., and Görlich, D. (2010). NES consensus redefined by structures of PKI-type and Rev-type nuclear export signals bound to CRM1. Nat Struct Mol Biol 17, 1367–1376. 10.1038/nsmb.1931.

39. Clarke, B.P., Angelos, A.E., Mei, M., Hill, P.S., Xie, Y., and Ren, Y. (2023). Cryo-EM structure of the CBC-ALYREF complex (elife) 10.7554/eLife.91432.1.

40. Ding, S., Wu, X., Li, G., Han, M., Zhuang, Y., and Xu, T. (2005). Efficient Transposition of the piggyBac (PB) Transposon in Mammalian Cells and Mice. Cell 122, 473–483. 10.1016/j.cell.2005.07.013.

41. Worch, R., Niedzwiecka, A., Stepinski, J., Mazza, C., Jankowska-Anyszka, M., Darzynkiewicz, E., Cusack, S., and Stolarski, R. (2005). Specificity of recognition of mRNA 5′ cap by human nuclear cap-binding complex. RNA 11, 1355–1363. 10.1261/rna.2850705.

42. Fornerod, M., Ohno, M., Yoshida, M., and Mattaj, I.W. (1997). CRM1 Is an Export Receptor for Leucine-Rich Nuclear Export Signals. Cell 90, 1051–1060. 10.1016/S0092-8674(00)80371-2.

43. Jumper, J., Evans, R., Pritzel, A., Green, T., Figurnov, M., Ronneberger, O., Tunyasuvunakool, K., Bates, R., Žídek, A., Potapenko, A., et al. (2021). Highly accurate protein structure prediction with AlphaFold. Nature 596, 583–589. 10.1038/s41586-021-03819-2.

44. Bradley, D., Garand, C., Belda, H., Gagnon-Arsenault, I., Treeck, M., Elowe, S., and Landry, C.R. (2024). The substrate quality of CK2 target sites has a determinant role on their function and evolution. Cell Systems 15, 544–562.e8. 10.1016/j.cels.2024.05.005.

45. St-Denis, N., Gabriel, M., Turowec, J.P., Gloor, G.B., Li, S.S.-C., Gingras, A.-C., and Litchfield, D.W. (2015). Systematic investigation of hierarchical phosphorylation by protein kinase CK2. Journal of Proteomics 118, 49–62. 10.1016/j.jprot.2014.10.020.

46. Abramson, J., Adler, J., Dunger, J., Evans, R., Green, T., Pritzel, A., Ronneberger, O., Willmore, L., Ballard, A.J., Bambrick, J., et al. (2024). Accurate structure prediction of biomolecular interactions with AlphaFold 3. Nature 630, 493–500. 10.1038/s41586-024-07487-w.

47. Melko, M., Winczura, K., Rouvière, J.O., Oborská-Oplová, M., Andersen, P.K., and Heick Jensen, T. (2020). Mapping domains of ARS2 critical for its RNA decay capacity. Nucleic Acids Research 48, 6943–6953. 10.1093/nar/gkaa445.

48. Liu, J.F., Hawley, B.R., Nicholson, L., and Jaffrey, S.R. (2024). Decoding m6Am by simultaneous transcription-start mapping and methylation quantification. BioRxiv, https://www.biorxiv.org/content/10.1101/2024.10.16.618717v1.

49. Kruse, T., Garvanska, D.H., Varga, J.K., Garland, W., McEwan, B.C., Hein, J.B., Weisser, M.B., Benavides-Puy, I., Chan, C.B., Sotelo-Parrilla, P., et al. (2024). Substrate recognition principles for the PP2A-B55 protein phosphatase. Sci. Adv. 10, eadp5491. 10.1126/sciadv.adp5491.

50. Punjani, A., Rubinstein, J.L., Fleet, D.J., and Brubaker, M.A. (2017). cryoSPARC: algorithms for rapid unsupervised cryo-EM structure determination. Nat Methods 14, 290–296. 10.1038/nmeth.4169.

51. Emsley, P., and Cowtan, K. (2004). *Coot* : model-building tools for molecular graphics. Acta Crystallogr D Biol Crystallogr 60, 2126–2132. 10.1107/S0907444904019158.

52. Afonine, P.V., Poon, B.K., Read, R.J., Sobolev, O.V., Terwilliger, T.C., Urzhumtsev, A., and Adams, P.D. (2018). Real-space refinement in *PHENIX* for cryo-EM and crystallography. Acta Crystallogr D Struct Biol 74, 531–544. 10.1107/S2059798318006551.

53. Pettersen, E.F., Goddard, T.D., Huang, C.C., Meng, E.C., Couch, G.S., Croll, T.I., Morris, J.H., and Ferrin, T.E. (2021). UCSF ChimeraX: Structure visualization for researchers, educators, and developers. Protein Sci 30, 70–82. 10.1002/pro.3943.

54. Kandiah, E., Giraud, T., De Maria Antolinos, A., Dobias, F., Effantin, G., Flot, D., Hons, M., Schoehn, G., Susini, J., Svensson, O., et al. (2019). CM01: a facility for cryo-electron microscopy at the European Synchrotron. Acta Crystallogr D Struct Biol 75, 528–535. 10.1107/S2059798319006880.

